# Coevolving Residues Distant from the Ligand Binding Site are Involved in GAF Domain Function

**DOI:** 10.1101/2024.08.07.605472

**Authors:** Wesam S. Ahmed, Anupriya M Geethakumari, Asfia Sultana, Anmol Tiwari, Tausif Altamash, Najla Arshad, Sandhya S Visweswariah, Kabir H Biswas

## Abstract

Ligand binding to GAF domains regulates the activity of associated catalytic domains in a wide variety of proteins. For instance, cGMP binding to the GAFa domain of phosphodiesterase 5 (PDE5) activates the cGMP-hydrolyzing catalytic domain in the protein. However, the residues involved and the mechanism of GAF domain function are not entirely clear. Here, combining computational and experimental analysis, we show that two highly coevolving residues distant from the ligand binding site play a critical role in GAF domain allostery. Specifically, Statistical Coupling Analysis (SCA) of GAF domain sequences revealed the highest coevolution score for residues L267 and F295. Molecular dynamics (MD) simulations of both apo and holo forms of the wild type and mutant (L267A and F295A) PDE5 GAFa domains revealed significant alterations in structural dynamics and interaction with cGMP. Incorporation of the mutations in a Bioluminescence Resonance Energy Transfer (BRET)-based biosensor, which reports a ligand-induced conformational change, revealed a change in the conformation of the GAF domain and an increase in the EC_50_ of cGMP-induced conformational change. Similar results were obtained regarding cGMP-induced conformational change in the full-length PDE5 and in the fluorescence of the GAF domain fluorescent protein, miRFP670nano3. Finally, structural analysis of conformers observed in MD simulations revealed a possible mechanism underlying the impact of mutations of these two coevolving residues in the PDE5 GAFa domain. Our results provide insight into the role of distant, coevolving residues in GAF domain allostery, and may aid in understanding evolution of allostery in proteins.

## Introduction

The activity of natural proteins, such as enzymes, is controlled by associated regulatory domains, which, in many cases, is achieved by binding a small molecule ligand and relaying associated structural changes to the catalytic domains. While many such regulatory domains are known, GAF domains (c**G**MP-specific PDEs, bacterial **A**denylyl cyclases, and bacterial **F**hLA transcriptional regulators) represent one of the largest family of regulatory domains [1], and are conserved in organisms ranging from archaea to chordates including humans [2]. Indeed, GAF domains play a crucial role in regulating protein function that impacts a range of biological processes, including gene expression regulation in bacteria [3], light-detection and signaling in plant and cyanobacteria [4], ethylene detection and signaling in plants [5], nitrogen fixation in bacteria [6], cAMP binding feedback control in cyanobacterial adenylyl cyclase [7], and the two-component sensor histidine kinase in both bacteria and plants [8, 9], as well as some of the mammalian cyclic nucleotide phosphodiesterases (PDEs) [1, 3, 10].

One of the well-studied GAF domains is the cGMP-binding GAFa domain in PDE5, which regulates cGMP levels in several human tissues, including vascular smooth muscle cells, lung, brain, kidney, cardiac myocytes, gastrointestinal tissue, platelets, and penile corpus cavernosum.[11, 12] The N-terminally located GAFa domain in PDE5 binds cGMP with a relatively high affinity and specificity and is thought to induce a conformational change in the domain that is transduced to the C-terminally located, cGMP-hydrolyzing catalytic domain, thereby activating it and increasing cGMP hydrolysis.[13, 14] Previous studies have shown that the apo GAF domain has high flexibility and that the conformational changes accompanyingcNMP binding stabilizes secondary structural elements in the domain.[15, 16] This converts the GAF domain from an “open” to a “closed” conformation wherein the cNMP is deeply buried inside the ligand binding site. Overall, there is still a limited understanding of the complexity of this allosteric communication induced by cNMP binding to the GAF domain, including the identity of amino acid residues that are involved in the process.

In the current study, we combined protein sequence-based coevolutionary analysis, molecular dynamics (MD) simulations, and biophysical assays, to better understand ligand binding and allosteric regulation in GAF domains (Fig. 1). Specifically, we performed Statistical Coupling Analysis (SCA) [17, 18] of GAF domain sequences to identify coevolving residue positions in the domain that may play a role in ligand binding and regulatory function and identified nine residue positions that showed high statistical coupling scores. Mapping these positions on the GAFa domain of PDE5 revealed two residues, namely L267 and F295, that showed the highest statistical coupling score and were located distant from the cGMP binding site. MD simulation analysis of the wild-type (WT) and mutant (both apo and cGMP-bound, holo forms) PDE5 GAFa domains revealed distinct changes in the structural dynamics of L267A and F295A mutants. Functional characterization using Bioluminescence Resonance Energy Transfer (BRET)-based biosensors revealed structural change in the basal state and an increase in the *EC*_50_ values of cGMP-induced conformational change in the mutant GAF domains (both isolated GAFa domain as well as in the full-length PDE5 protein). Moreover, genomic database and structure-based analysis predicted that variations in these two positions in PDE5A gene were deleterious. Importantly, equivalent mutations in the fluorescent GAF domain, miRFP670nano3, resulted in a decrease in the fluorescence of the protein indicative of a conserved role of these two distant, coevolving residues in GAF domain function.

**Fig. 1.**
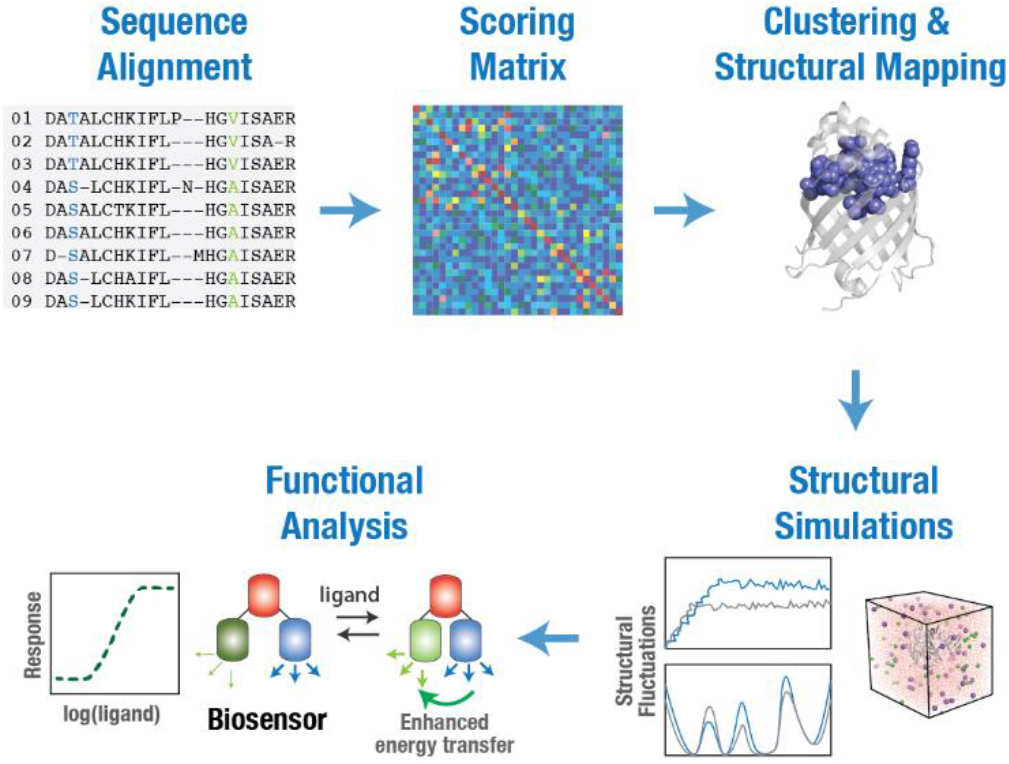
Coevolving residue positions and their role in GAF domain allostery. Schematic showing an approach to determine coevolving residues and delineating their role in the allosteric regulation of GAF domains. GAF domain sequences were aligned, and statistical coupling analysis (SCA) was performed to detect coevolving residue positions. The scoring matrix obtained from SCA was then used for clustering to determine the cluster of coevolving residues with highest scores, which were then mapped on to a GAF domain structure. MD simulations followed by biophysical assays with biosensors were then used to determine the role of individual coevolving residue positions in the allosteric regulation of GAF domains.

## Materials and Method

### Identification of coevolving residues in GAF domains

To perform SCA [19, 20], GAF domain protein sequence alignment was obtained from the Pfam database [21] and phylogenetically related sequences were removed using JalView [22] by applying a redundancy threshold value of 95%. The dataset was further refined by removing all positions with more than 20% gaps using available MATLAB scripts. SCA was performed using the algorithm described previously [18]. Hierarchical clustering analysis of the statistical coupling scores was performed by constructing a dendrogram of the correlation matrix using MATLAB. Cluster of residue positions with the highest statistical coupling scores were mapped on the PDE5 GAFa domain structure (PDB: 2K31[23]).

### MD simulation and trajectory analysis

MD simulations were performed using the available structure of apo (PDB: 3MF0) [53] or modeled structure of the holo PDE5 GAFa domain. Homology modeling was performed using the mouse holo GAFa domain (PDB: 2K31; amino acid residues 154-302) [23] (Supporting Figure 1) using MODELLER (10.1 release, Mar. 18, 2021) [24]. PROCHECK was used to assess the stereochemical quality of the model (Supporting Figure 2) and the Cα atom root-mean-square deviation (RMSD) between the model and the template was found to be 0.6 Å. Structures of the mutants were generated using the PyMOL mutagenesis tool. Parameter and topology files for the MD simulations were generated using the CHARMM-GUI server [25] and simulations were performed using the NAMD v2.13 software [26] and CHARMM36 force field [27] as described previously [28-32]. Structures were solvated using a TIP3P [40] cubic water box with a 10 Å minimum distance between the edge of the box and any of the biomolecular system atoms, and NaCl was added at a final concentration of 0.15 M with periodic boundary conditions applied. Before performing MD simulation production runs, energy minimization was first performed for 1000 timesteps, followed by a thermalization step where the systems were slowly heated to 310 K for 0.25 ns using a temperature ramp that raises the temperature at 1 K increment. Temperature was then maintained at 310 K using Langevin temperature control and at pressure of 1.0 atm using Nose-Hoover Langevin piston control. A constrained equilibration step of 1 ns was then performed where protein backbone atoms were constrained using a harmonic potential. Finally, 1000 ns MD simulation production runs were performed in triplicates for the WT and mutant systems. A 2 fs timestep of integration was chosen for all simulations where short-range non-bonded interactions were handled at 12 Å cut-off with 10 Å switching distance, while Particle-mesh Ewald (PME) scheme was used to handle long-range electrostatic interactions at 1 Å PME grid spacing. Analysis of the trajectories was performed using the available tools in the VMD software [33]. Dynamic cross-correlation (DCC) analysis was carried out as described previously [28, 34] using the Bio3D package in R software [35]. Binding free energy changes were estimated utilizing the MM-PBSA method [36] through the CaFE 1.0 plugin [37] in the VMD software [33] using a Tcl script (Supporting Text 6).

### Plasmid design and mutagenesis

The design of the BRET^2^-based GAFa and full-length PDE5 (A2 isoform) biosensors was described previously [38, 39]. Briefly, in both biosensors, the GAFa domain (I159-N316) or the full-length PDE5A2, were flanked by the green fluorescent protein 2 (GFP^2^) at the N-terminal as the fluorescent energy acceptor, and by the *Renilla* luciferase (RLuc) at the C-terminal as the bioluminescent energy donor. In vitro, site-directed mutagenesis of the GAFa-based biosensor was performed using two mutagenic primers (5’ G ACA CAA AGC ATT GCC TGT ATG CCA ATT AAG 3’) and (5’ CA GGA AAC GGT GGT ACC GCT ACT GAA AAA GAT G 3’) to generate the L267A and F295A mutations in the GAFa domain, respectively, in a pBKS-GAFa plasmid. Mutations were confirmed through sequencing. The L267A and F295A mutant GAFa domain (residues I159-N316) sequences were then inserted into pGFP^2^-MCS-RLuc plasmid through restriction digestion using *Bgl*II and *Hind*III restriction enzymes and subsequent ligation to generate the GAFa-based mutant biosensor constructs (Supporting Text 1) [38]. Mutations in the full-length PDE5A2 biosensor (pGPF^2^-PDE5A2-RLuc) [40] were generated by GenScript (GenScript, Singapore) (Supporting Text 2).

To generate the mNeonGreen-GAFa-NanoLuc (mNG-GAFa-NLuc) biosensor construct, the GAFa sequence was synthesized and inserted into the pmNG-Rigidx3-NLuc plasmid using *Kpn*I-*Not*I restriction sites to generate the pmNG-GAFa-NLuc plasmid. For generating the miRFP670nano3-picALuc(E50A) construct, the miRFPnano3 gene fragment was synthesized and inserted into the pcDNA3.1-mSca-M^pro^-Nter-picALuc(E50A)[41] using the *Eco*RI-*Bam*HI restriction sites to generate the pcDNA3.1-miRFP670nano3-picALuc(E50A) plasmid. L267A and F295A mutants of mNG-GAFa-NLuc, and L100A and W125A mutants of miRFP670nano3-picALuc(E50A) were then generated in their respective plasmids (GenScript, Singapore; Supporting Text 3, 4).

### Cell culture, transfection, and western blot analysis

The HEK293T cells were maintained in Dulbecco’s Modified Eagle’s Medium (DMEM) with 10% fetal calf serum (FCS), 120 mg/L penicillin and 270 mg/L streptomycin at 37 °C in a humidified incubator with an atmosphere of 5% CO_2_. The cells were transfected with plasmids expressing the protein (WT or mutants) using either polyethyleneimine (PEI; Sigma-Aldrich, USA) or Lipofectamine 2000 (ThermoFisher Scientific, USA) transfection reagents.

For western blot analysis, lysates prepared from HEK293T cells expressing the biosensor constructs were subjected to SDS-PAGE, and proteins were transferred to nitrocellulose membranes. Blots were probed using an anti-GFP antibody for GAFa and full-length PDE5 sensors (mouse IgG2a κ mAb, Santa Cruz Biotechnology, USA – SC-9996; 1:500 dilution), an anti-FLAG tag antibody for miRFP670nano3 constructs (mouse IgG2a mAB, Sino Biological, China – 109143-MM13; 1:10000 dilution), or an anti-α-tubulin antibody for α-tubulin (mouse IgG2b mAB, Proteintech, USA – 66031-1; 1:200000). All blots were probed using the same secondary antibody (anti-Mouse IgG:HRP Donkey pAb; ECM Biosciences, USA – MS3001; 1:20000 dilution) and developed using a chemiluminescent substrate in a ChemiDoc Imaging System (BIO-RAD, Germany).

### In vitro BRET assays

In vitro BRET assays were performed using cell lysates obtained from HEK293T cells transfected with plasmids expressing the GAFa domain-based (GFP^2^-GAFa(I159-N316)-RLuc) or the full-length PDE5 (GFP^2^-PDE5A2-RLuc) WT or mutant biosensor constructs [7, 29, 38, 40, 42]. Cells were washed in chilled Dulbecco’s phosphate-buffered saline (DPBS) after 48 h of transfection, harvested using DPBS containing 2 mM EDTA, and lysed by sonication in a lysis buffer containing 50 mM HEPES (pH 7.5), 100 mM NaCl, 2 mM ethylenediaminetetraacetic acid (EDTA), 1 mM dithiothreitol (DTT), 1× protease inhibitor cocktail (ThermoFisher Scientific, USA) and 10% glycerol [38, 43]. Sonicated lysates were centrifuged at 4 °C for 1 h at 18,400g, and supernatants were collected. Bioluminescence spectra of the WT or the mutant biosensors were acquired over wavelengths ranging between 385 to 665 nm utilizing a grating-based system with 15 nm integration step size, with each individual measurement acquired for 1 second using a Tecan SPARK® multimode microplate reader (Tecan, Switzerland). BRET was determined as a ratio of GFP^2^ and RLuc emission, either in the absence or in the presence of the indicated concentrations of cGMP, following a 30-minute incubation at 37 °C. Bioluminescence and BRET were measured following the addition of the RLuc substrates either coelenterazine 400a or Prolume purple (NanoLight Technology, Prolume Ltd., USA).

### Determination of thermal stability of the GAFa domain of PDE5

Thermal stability of the WT and mutant GAFa domains was determined by measuring BRET from cell lysates obtained from HEK293T cells transfected with plasmids expressing the WT (mNG-GAFa-NLuc) or mutant (mNG-GAFa(L267A)-NLuc or mNG-GAFa(F295A)-NLuc) biosensors.[44] Cell lysates were incubated at a range of temperatures for 10 min using a C1000 Touch Thermal Cycler (BIO-RAD, Germany), after which BRET was measured using GloMax® DSiscover plate reader (Promega, USA). Data were fit to a Boltzmann sigmoidal model to determine the melting temperature, *T*_m_ (temperature at which the biosensor shows half maximal BRET). Three independent experiments were performed in triplicates.

### Determination of miRFP670nano3 fluorescence

Plasmids expressing the WT [pcDNA3.1-miRFP670nano3-picALuc(E50A)] and mutants [pcDNA3.1-miRFP670nano3(L100A)-picALuc(E50A) or pcDNA3.1-miRFP670nano3(W125A)-picALuc(E50A)] were transfected into HEK293T cells. After 48 h of transfection, miRF670Pnano3 fluorescence was measured using GloMax® Discover plate reader (Promega, USA), utilizing a 627 nm excitation filter and a 660-720 emission filter, and picALuc bioluminescence intensity was determined after the addition of coelenterazine h utilizing Tecan SPARK® multimode microplate reader (Tecan, Switzerland). Five independent experiments were performed in triplicates.

### Data analysis and figure preparation

GraphPad Prism (version 8) and Microsoft Excel (2016) were used for data analysis and graphs preparation. Figures were assembled using either Inkscape or Adobe Illustrator software.

## Results

### Identification of coevolving residues in GAF domains using SCA

To determine coevolving residues and better understand the allosteric communication in the GAF domain, we used the statistical coupling analysis (SCA) coevolutionary analysis method followed by MD simulations and functional analysis to determine the role of the identified coevolving residues in GAF domain allostery (Fig. 1). SCA involves analysis of covariation between pairs of amino acid positions in multiple sequence alignments (MSA) to identify coevolving residues in a protein family.[17, 18]It specifically measures the extent to which the distribution of amino acids at one position (x) is altered by changes in the amino acid distribution at another position (y). Towards this, GAF domain sequence alignment obtained from the Pfam database [21] was refined for phylogenetic relatedness and gaps, and SCA was performed using previously described algorithms [18, 45] to generate a matrix of pairwise correlation scores for each residue in the GAF domain (Fig. 2A). Importantly, only ∼1% of the residue pairs showed correlation scores >0.25, and hierarchical clustering analysis revealed a cluster of nine highly coevolving residue positions (Supporting Figure 3), suggestive of their role in ligand binding and allostery in GAF domains.

**Fig. 2.**
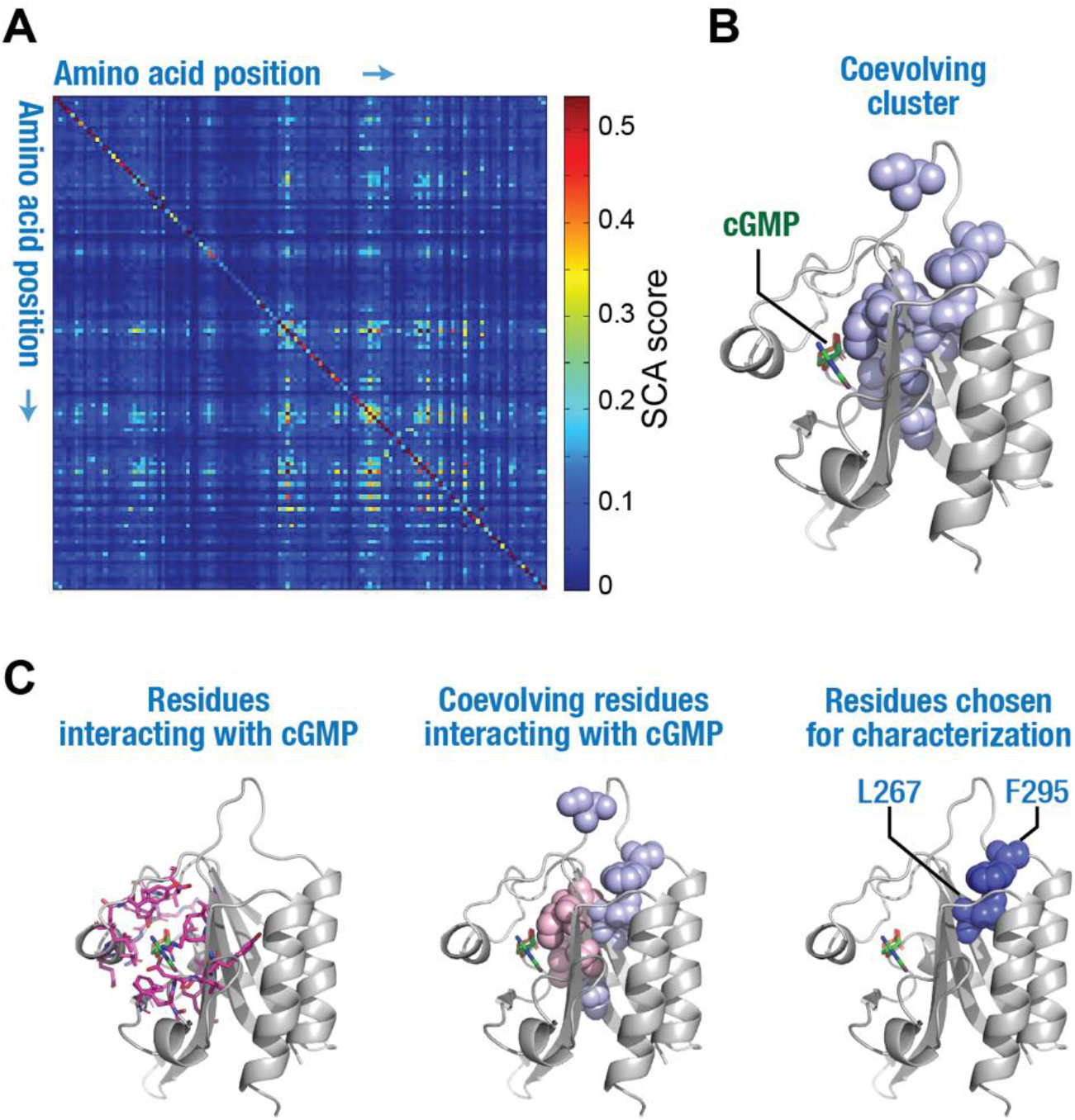
SCA reveals a cluster of coevolving residues in GAF domains. (A) Color-coded heat map showing pairwise scores of individual residue positions in GAF domains obtained from SCA. (B) Cartoon representation of the GAFa domain of PDE5 (PDB: 2K31[23]) showing the coevolving cluster of residues (spheres in light blue) in the GAF domains. (C) Cartoon representation of the GAFa domain of PDE5 showing residues (side chains as sticks in pink) within a 5 Å distance from cGMP (left panel), coevolving residues (spheres in light pink; rest of the coevolving residues in light blue) within 5 Å distance from cGMP (middle panel) and coevolving residues L267 and F295 selected for further functional characterization (dark blue spheres, right panel).

We mapped the cluster of coevolving residues on the cGMP-bound structure of the GAFa domain of PDE5 (amino acid residues 164-312; human PDE5A1 numbering; PDB: 2K31 [23]) (Fig. 2B). We chose to use the GAFa domain of PDE5 for the following reasons: first, unlike other GAF domains that are either not known to bind small molecules or whose function is still unknown, the PDE5A GAFa domain has a well-known function, which is to bind cGMP with high specificity and affinity, and allosterically activate the cGMP-hydrolyzing activity of PDE5 catalytic domain. The isolated PDE5 GAFa domain has been reported to show a thousand-fold higher affinity for cGMP compared to cAMP [23]. This selectivity is much higher compared to some other cGMP-binding GAF domains such as the cGMP-binding GAFb domain of PDE2A, which shows only about 20-fold higher affinity for cGMP [46]. Second, the three-dimensional structures of the isolated PDE5 GAFa domain both in the absence of cGMP (unliganded; apo) [15] and cGMP-bound (holo) [23] forms are available in the literature, which allows for robust structural comparison and subsequent computational investigations. Third, we have previously engineered BRET-based conformational biosensors for both the isolated GAFa domain as well as the full-length PDE5 that faithfully report cGMP-binding induced conformational changes in the proteins, which has enabled the characterization of several mutations with regards to GAF domain allostery [38, 40, 43].

In the human PDE5, the GAFa domain is composed of six antiparallel β sheets (β1-β6) and four α helices (α2-α5) with an arrangement of the secondary structural elements of αβββαβαββα [15]. Positional mapping of the nine coevolving residue positions (Fig. 2B) revealed that six residues (V230 located on helix α3, I266 and C268 located on sheet β5, V281, Q283, and I285 located on sheet β6; all residue numbers indicated are as per human PDE5A1 sequence) are located in (V230, I266, Q283, I285) or near (C268, V281) the ligand binding site in the GAFa domain (binding site residues were identified using a cutoff of 5 Å [47] distance from cGMP in the holo GAFa domain [23]). On the other hand, the remaining three residues (S289 in the loop between sheet β6 and helix α5, L267 in sheet β5, and F295 in the loop between sheet β6 and helix α5) are located distant from the binding site with no direct interaction with the cGMP (Fig. 2C, Supporting Movie 1). Interestingly, two of the distantly located residues (L267 and F295) had the highest coupling scores among all the nine coevolving residues identified from SCA. A closer inspection of these two residues revealed that they form a hydrophobic interaction (Supporting Movies 2, 3). Therefore, of the nine coevolving residues, we chose to perform further characterization of the L267 and F295 residue positions individually in the PDE5 GAFa domain.

### MD simulations reveal distinct changes in the structural dynamics of the PDE5 GAFa domain upon mutation of coevolving residues distant to the cGMP binding site

To evaluate the structural and functional impact of the two highly coevolving positions (L267 and F295) we individually generated L267A and F295A mutations in the apo and holo structural models of the GAFa domain. We then performed three independent, all-atom, explicit solvent, 1000 ns-long, MD simulation runs of the WT and mutant GAFa domains in both the apo and holo forms. MD simulation has found several applications, including probing the effect of single-point mutations on the structure of proteins [48] and the determination of ligand binding affinity [47]. A detailed analysis of the MD simulation trajectories revealed several distinct features in the apo and holo GAFa domain structural dynamics, in addition to several differences between the WT and the two mutant GAFa domains. Comparison of the WT apo and holo forms of the GAFa domain confirmed previous data that the apo form had higher flexibility to accommodate ligand binding and subsequent conformational changes. (Fig. 3A). Specifically, analysis of the apo and holo WT GAFa domain trajectories revealed a reduced root-mean-square deviation (RMSD) value in the cGMP-bound GAFa domain (4.5 ± 0. 9 and 3.4 ± 0.7 Å for the apo and holo structures, respectively). Interestingly, while the apo-form of the L267A mutant showed a decrease (4.2 ± 0.7 Å) in RMSD, the F295A mutant apo-form showed an increase (4.5 ± 0.9) as compared to the WT GAFa domain(4.9 ± 0.9 Å). Importantly, a decrease in the RMSD was observed for the holo L267A as compared to the apo L267A mutant (4.2 ± 0.7 and 3.9 ± 1.0 Å respectively) as seen with the WT GAFa domain. However, no decrease in RMSD was observed for the holo F295A mutant (4.9 ± 0.9 and 4.9 ± 1.1 Å for the apo and holo F295A mutant, respectively) (Fig. 3B).

**Fig. 3.**
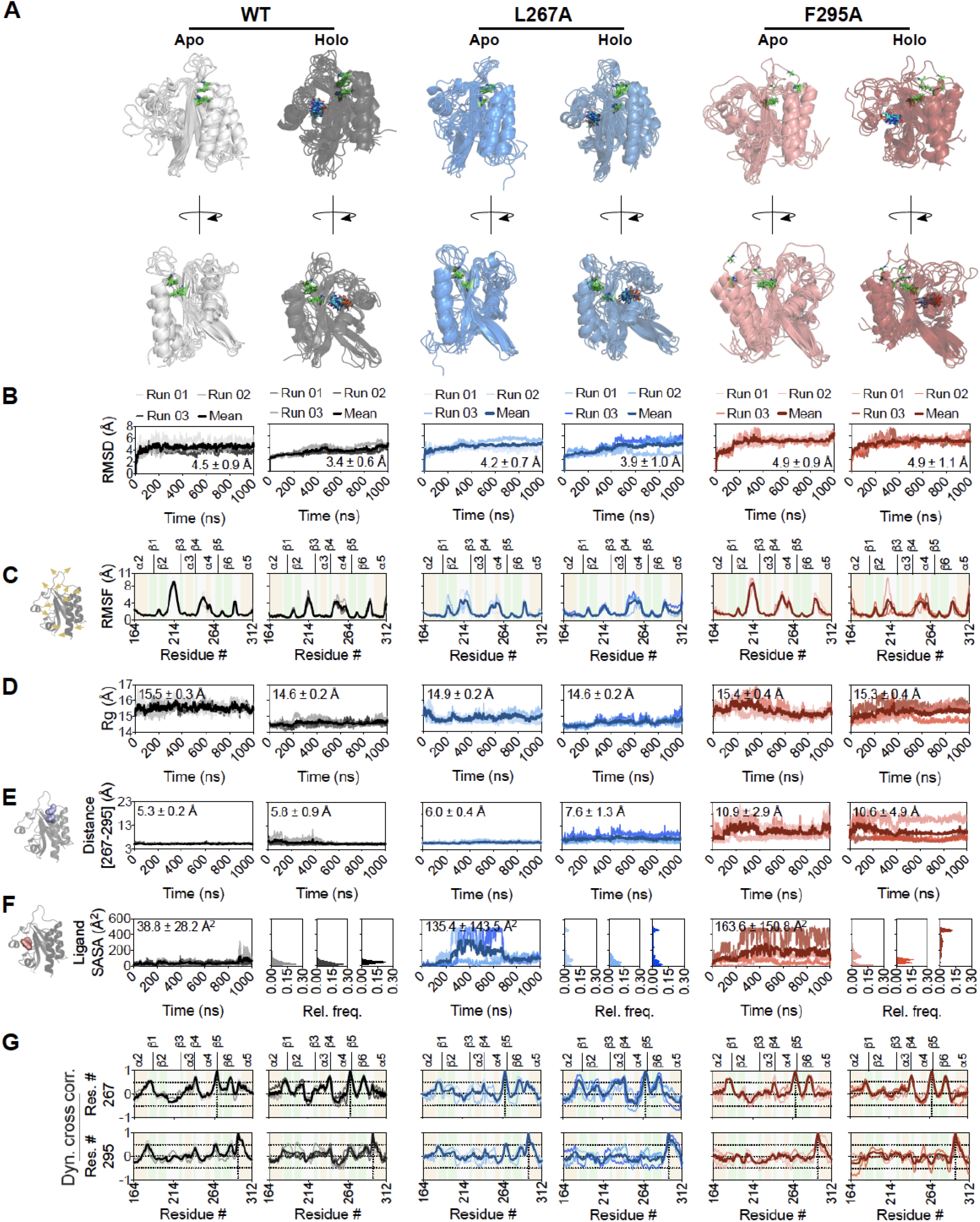
Mutation of distant, coevolving residue positions induces dynamic structural changes in the GAFa domain of PDE5. (A) Cartoon representation of GAFa domain (WT, left panel; L267A mutant, middle panel; and F295A mutant, right panel) dynamics across the 1000 ns simulation runs captured using a composite image of snapshots that are 200 ns apart. The ligand is shown as blue carbon sticks, and positions 267 and 295 are shown as green carbon sticks in each complex. (B) Graphs showing root mean squared deviation (RMSD) measurements of the apo and holo GAFa domains (WT and mutants) across the 1000 ns MD simulation runs. (C) Schematic (left panel) and graphs showing root mean square fluctuation (RMSF) measurements of the apo and holo GAFa domains (WT and mutants) across the 1000 ns MD simulation runs (right panel). (D) Graphs showing radius of gyration (Rg) of the apo and holo GAFa domains (WT and mutants) across the 1000 ns MD simulation runs. (E) Schematic highlighting residues L267 and F295 (left panel) and graphs showing center-of-mass distance between amino acid positions 267 and 295 in the apo and holo GAFa domains (WT and mutants; right panel). Note how the hydrophobic interaction is disrupted in the F295A mutant domain as is evident from the increased distance between the two positions, as opposed to the WT and L267A mutant GAFa domains that maintained the distance across the 1000 ns MD simulation runs. (F) Schematic highlighting ligand (cGMP) surface (left panel) and graph showing solvent accessible surface area (SASA) of cGMP across the 1000 ns MD simulation runs of the holo GAFa domain (WT and mutants; right panel). Outsets, frequency distribution of ligand SASA across the 1000 ns MD simulation runs of each complex. (G) Graphs showing dynamic cross-correlation (DCC) values of residue positions 267 (top panel) and 295 (bottom panel) in the apo and holo GAFa domain (WT and mutants) obtained from the 1000 ns MD simulation runs. Vertical dotted lines indicate the residue 267 in graphs in the top panel and residue 295 in graphs in the bottom panel, while horizontal dotted lines indicate DCC values of 0.5 and −0.5.

Inspecting the structural fluctuation of individual amino acid residues through root-mean-square fluctuation (RMSF) analysis showed that residues forming the loop regions (β1/β2, β2/β3, β4/α4, β5/β6, and β6/α5) of the domain exhibited higher fluctuations in both the apo and holo forms of the WT and mutant domains compared to other residues (Fig. 3C). Comparison of the apo and holo WT GAFa domain revealed a notable decrease in the fluctuation of the β2/β3 loop, which flanks the cGMP binding site, in the holo form. Interestingly, compared to the apo WT, the decrease in the fluctuation of this loop was also observed in the apo L267A mutant, but not the apo F295A mutant. In this regard, the holo L267A mutant showed a minute decrease in the fluctuation of this region compared to the apo L267A mutant, contrary to the WT and F295A mutant domains, where the fluctuation of this region was notably decreased in the holo form of each domain compared to its apo form (Fig 3C).

We then performed radius of gyration (*R*g) analysis of the trajectories to determine the changes in the overall structure of the proteins (Fig. 3D). This revealed a decrease in the *R*g of the holo WT GAFa domain in comparison to the apo (15.5 ± 0.3 and 14.6 ± 0.2 Å for the apo and holo structures, respectively) (Fig. 3D). Additionally, comparing the apo WT and mutant GAFa domains revealed a notable decrease in the *R*g in the apo L267A mutant compared to the apo WT, while a similar decrease was not observed for the apo F295A mutant (15.5 ± 0.3, 14.9 ± 0.2 and 15.4 ± 0.4 Å for the apo WT and the L267A and F295A mutant domains, respectively). However, in the holo form, the F295A mutant showed an increase in the *R*g compared to holo WT, while this increase was not reflected in the holo L267A mutant (14.6 ± 0.2, 14.9 ± 0.2 and 15.3 ± 0.4 Å for the holo WT, L267A and F295A mutant domains, respectively) (Fig. 3D).

We then determined the distance (center-of-mass) between the highly coevolving residue positions 267 and 295 from the simulation trajectories to monitor their interaction and how their interaction is altered, if at all, by the mutations in these positions (Fig. 3E). This analysis revealed a close positioning of the two residues in the apo WT GAFa domain throughout the 1000 ns-long MD simulation trajectories (Fig. 3E). A marginal increase in the distance was observed in the holo WT GAFa domain trajectories (5.3 ± 0.2 and 5.8 ± 0.9 Å for the apo and holo GAFa domain trajectories, respectively) (Fig. 3E). The L267A mutation resulted in an increase in the distance in the apo state, as compared to the WT GAFa domain (5.3 ± 0.2 and 6.0 ± 0.4 Å for the WT and the L267A mutant GAFa domain, respectively). The distance increased further in the holo L267A GAFa domain (6.0 ± 0.4 and 7.6 ± 1.3 Å for the apo and holo L267A mutant GAFa domain, respectively). Interestingly, the distance was found to be much larger, with greater fluctuation, in the case of the F295A mutant GAFa domain (10.9 ± 2.9 and 10.6 ± 4.9 Å for the apo and holo F295A mutant GAFa domain, respectively), in comparison to both the WT and the L267A mutant GAFa domain (Fig. 3E), suggesting a loss of the interaction between the two residues in the F295A mutant GAFa domain.

Overall, these results show that the apo WT GAFa domain is more dynamic and assumes a more compact form in the holo state. Indeed, it has been suggested that cGMP binding to the GAFa domain of PDE5 converts the domain from an ‘open’ state conformation into a more compact structure following ligand binding [23]. Similar changes were reported for an isolated cAMP-binding GAF domain [7]. Additionally, the L267A mutant appears to convert the apo GAFa domain into a more compact structure, which is accompanied by a decrease in the fluctuation of the β2/β3 loop, which could negatively impact cGMP-binding where the flexibility of the apo domain is an important feature. On the other hand, the F295A mutant disrupts the hydrophobic interaction between these two positions and other positions in the same region (Fig. 3A,E). This results in an increase in the dynamics and a decrease in the compactness of the domain most noticeable in the holo form, which is likely to affect the ligand binding and the subsequent cGMP-binding induced conformational change.

To investigate how these structural changes in the WT and L267A and F295A mutant GAFa domain affect cGMP inside the ligand binding site, we determined the solvent-accessible surface area (SASA) of cGMP from the holo GAFa domain MD simulation trajectories. This analysis revealed large increases in the cGMP SASA in both the L267A (38.8 ± 28.2 Å) and F295A (135.4 ± 143.5 Å) mutant GAFa domains, as compared to the WT GAFa domain 163.6 ± 150.8 Å) (Fig. 3F). These results suggest that the ligand is less buried inside the binding pocket in the case of the mutant holo GAF domains compared to the WT.

To understand how ligand binding may be affected by these mutations, we calculated the interaction energy (van der Waal’s and electrostatic) and hydrogen bonding (H-bonding) between cGMP and GAFa domain using the three independent MD simulation trajectories of each of the GAFa domains. This analysis revealed a decrease in both types of interactions in the mutants (Supporting Figure 5), suggesting that ligand interaction with the binding site of GAFa domain is negatively affected due to the mutations. Overall, these results suggest that L267A and F295A mutations result in alterations in the structure and dynamics of the GAFa domain that are likely to affect cGMP-binding induced conformational change.

Finally, we performed dynamic cross-correlation (DCC) analysis of the MD simulation trajectories of the WT, L267A and F295A mutant GAFa domains to understand the changes in the pair-wise correlated motions of residues in the proteins (Supporting Figure 4). A comparison of the DCC plots of the apo and holo WT GAFa domain revealed an increase in the DCC values between residue positions 244-264 (which form the α4 region), and positions 184-210 (which form the β1/β2 region) and 264-290 (which form the β4/β5 region) (Supporting Figure 4). The dynamically cross-correlated motions between these residue positions were further increased in the holo L267A GAFa domain compared to the holo WT GAFa domain, while such increases were not apparent in the holo F295A mutant GAFa domain. On the other hand, the apo L267A mutant GAFa domain appeared to disrupt the dynamically cross-correlated motions that were originally observed in the apo WT GAFa domain. On the contrary, the F295A mutant GAFa domain did not appear to show any notable impact on these motions (Supporting Figure 4).

We then focused our attention on the DCC of the residue positions 267 and 295 against all other residues in the proteins (Fig. 3G). While both positions 267 and 295 showed highly correlated motions (equal to or above 0.5) with residues in the α2/β1 loop, sheets β4 (which flanks the cGMP binding site) and β6 in the apo WT GAFa domain trajectories, the position 295 showed additional correlated motions with residues on its C-terminal side in the helix α5 (Fig. 3G). The residue position 267 showed increased correlated motions with residues in the β2/β3 loop while residue position 295 showed a decrease in correlated motions with residues in the α2/β3 loop in the holo, as compared to the apo, WT GAFa domain. A decrease in the correlated motions of residues positions 267 and 295 with residues in sheets β4 was observed in the apo L267A and F295A mutant, as compared to the WT, GAFa domain. Further, no apparently large differences were observed for the correlated motions of residue positions 267 and 295 in both the holo L267A and F295A mutant, as compared to the WT, GAFa domain (Fig. 3G). Overall, these results indicate a decrease in the dynamically cross-correlated motions between the two coevolving positions and specific regions in the L267A and F295A mutants, as compared to the WT GAFa domain.

### L267A and F295A mutations result in an increase in the EC_50_ of cGMP-induced conformational change in PDE5 GAFa domain

Having observed alterations in the structural dynamics in the GAFa domain of PDE5 upon L267A and F295A mutations using MD simulations, we then proceeded to determine the impact of these mutations experimentally. In this regard, structural studies on the GAFa domain of PDE5 have revealed the conformational changes in the protein upon cGMP binding (Fig. 4A). Broadly, these changes include a close juxtapositioning of the sheet β3 and helix α4 leading to a ‘closing’ of the cGMP binding site, an ‘inward’ movement of sheet β1 and β2 towards the cGMP binding site and an ‘outward’ movement of the loop β2/β3 (Fig. 4A, left panel). Additionally, cGMP binding is associated with a close juxtapositioning of the helices α2 and α5 at the N- and C-termini of the GAFa domain, likely associated with the transduction of allosteric signal from the GAFa domain to the catalytic domain in PDE5 (Fig. 4A, right panel). We decided to take advantage of the latter to monitor cGMP-induced conformational change in the GAFa through an increase in the BRET efficiency of the GAFa domain biosensor that we have described previously [49] (Fig. 4B). BRET is a biophysical technique that involves resonance energy transfer between a bioluminescent donor and a fluorescent acceptor, the extent of which is dependent on the spectral overlap, distance, and relative orientation between the two reporters. In this case, the biosensor consists of an isolated GAFa domain of PDE5 sandwiched between GFP^2^ at the N-terminal side (BRET acceptor) and RLuc at the C-terminal side (BRET donor). GFP^2^ is a brighter variant of GFP with a F64L mutation and has an emission spectra similar to that of GFP but with significantly blue-shifted excitation spectra with a peak of 396 nm (Patent no. US 6.12,188 B1) [50], leading to a substantial spectral overlap of RLuc emission with GFP^2^ excitation as well as a large separation of RLuc and GFP^2^ emissions [38, 40, 43].

**Fig. 4.**
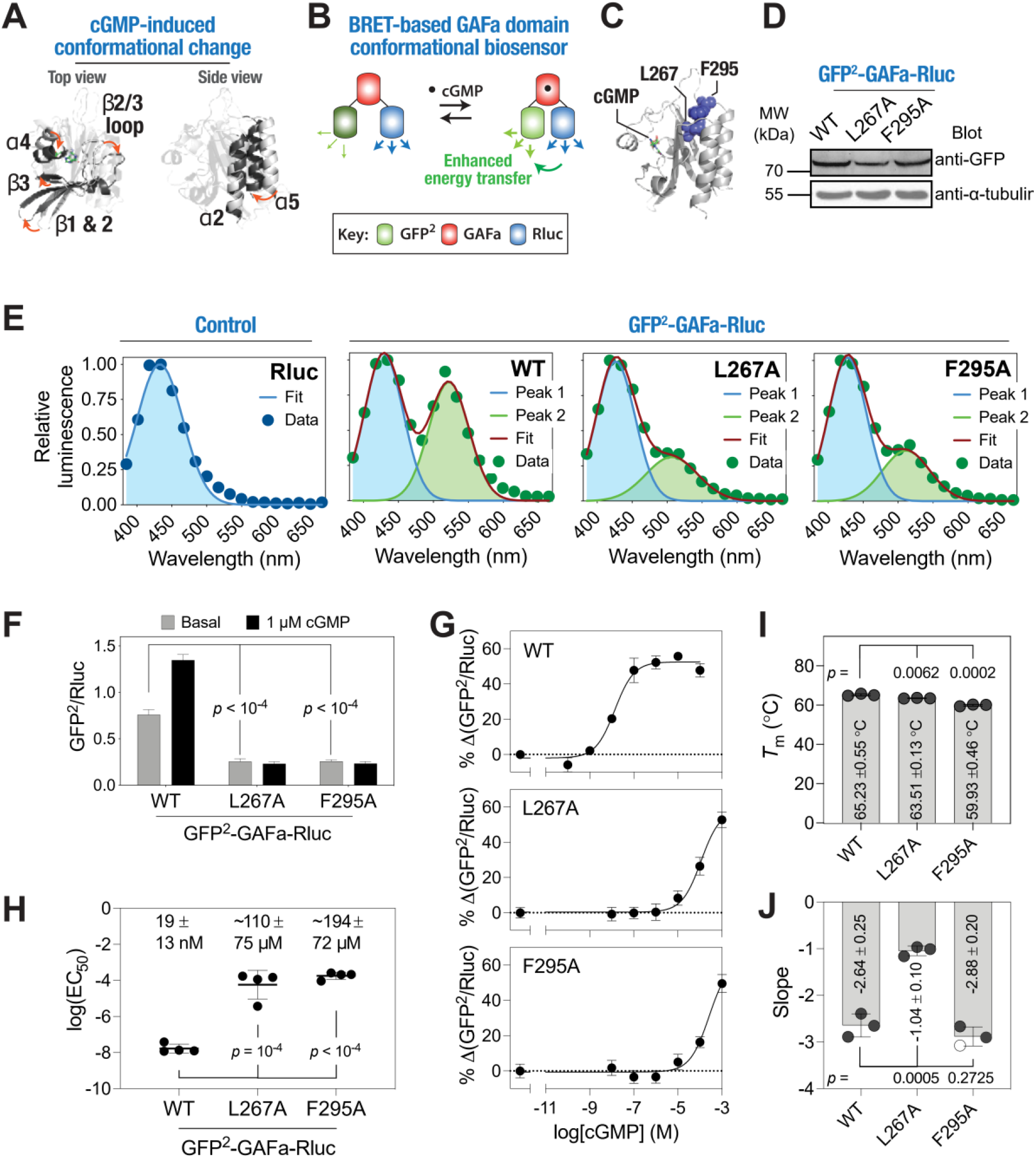
Mutation of the distant, coevolving residues alters cGMP-induced allosteric conformational change in the isolated GAFa domain of PDE5. (A) Cartoon representation of apo (PDB: 3MF0[15]) and holo (PBD: 2K31[23]) PDE5 GAFa domain structures aligned using all Cα-atoms (left panel) and using Cα-atom of α2 helix (right panel) highlighting structural changes (apo: gray, holo: black) induced upon cGMP binding. (B) Schematic showing the BRET^2^-based GAFa domain conformational biosensor that shows increased energy transfer upon cGMP binding likely due to the structural juxta positioning of the helix α5 against helix α2, and thus, bringing the BRET donor and acceptor closer. (C) Cartoon representation of the PDE5 GAFa domain highlighting the distant, coevolving residues L267 and F295 (spheres in blue) that were mutated to A. The ligand, cGMP, is shown in the stick representation. (D) Western blot analysis of the cell lysates prepared from HEK293T cells transfected with either the WT or mutant GAFa domain biosensor plasmids probed using an anti-GFP antibody showing the expression of biosensor constructs. Whole blot image used for generating the figure is shown in Supporting Figure 7. (E) Graphs showing bioluminescence spectra obtained from lysates prepared from cells expressing either RLuc alone (control) or the indicated GAFa biosensor constructs (containing the WT or the mutant PDE5 GAFa domains sandwiched between GFP^2^ and RLuc proteins). Note the appearance of a second peak in the WT GAFa domain biosensor, indicating energy transfer from RLuc (donor) to GFP^2^ (acceptor), while this peak is reduced in both L267A and F295A mutant GAFa domain biosensors. Data shown are representative of multiple experiments and were fit using either a single (RLuc alone) or two Gaussian distributions (GAFa domain biosensors). (F) Graph showing BRET values (ratio of GFP^2^ and RLuc emissions) of the WT, L267A, and F295A mutant GAFa domain biosensors in the absence and presence of cGMP (1 μM). Note the significant reduction in the basal BRET values of the L267A and F295A mutant biosensors compared to the WT biosensor. (G) Graphs showing percentage increases in BRET for the WT, L267A, and F295A mutant GAFa domain biosensors with the indicated cGMP concentrations. Note the shift in the cGMP dose-response curves of the L267A and F295A mutant GAFa domain biosensors in comparison to the WT GAFa domain. Data shown are the mean ± standard deviation (S.D.; error bars) from a representative experiment, with experiments performed four times. (H) Graph showing log(*EC*_50_) values of cGMP-induced conformational change in the WT, L267A, and F295A mutant GAFa domain biosensor. Data shown are mean ± S.D. obtained from four independent experiments. Inset, values on top indicate the cGMP *EC*_50_ values (mean ± S.D.) of the respective GAFa domain biosensors. (I,J) Graph showing melting temperatures (*T*_m_) and slope of the melting temperature curves of the WT and L267A and F295A mutant GAFa domain biosensors. Data shown are mean ± S.D. obtained from three independent melting temperature curves (curves are provided in Supporting Figure 8). *p*-values shown were obtained from Student’s t-tests (unpaired, equal variance) performed for the L267A and F295A mutants against the WT GAFa domain. Numbers shown in the graphs are mean ± S.D. obtained from three independent experiments. For all tests, a *p*-value of <0.05 was considered statistically significant.

We generated L267A and F295A mutations individually in the BRET-based GAFa domain biosensor plasmid [38] (Fig. 4C) and confirmed the expression of the biosensors through western blot analysis (Fig. 4D) Determination of bioluminescence spectra revealed a second peak in the case of the WT GAFa domain biosensor that was not observed in the case of the RLuc alone control (Fig. 4E), indicating a significant resonance energy transfer from RLuc to GFP^2^. On the other hand, the mutant GAFa domain biosensors showed a reduced second peak, indicating a decrease in the energy transfer from RLuc to GFP^2^. Further, basal BRET ratio measurements showed statistically significant decrease in the mutants (0.76 ± 0.05, 0.25 ± 0.03, and 0.26 ± 0.02 for WT, L267A, and F295A, respectively; *p* < 10^−4^ for WT vs L267A and *p* < 10^−4^ for WT vs F295A), suggesting a conformational change in the L267A and F295A mutant GAFa domains.

Further, we incubated the biosensors with 1 µM cGMP, a relatively high concentration as the PDE5 GAFa domain binds cGMP with nanomolar affinity [38], and measured BRET. In agreement with our previous report [38], the WT GAFa domain biosensor showed an increase in BRET in the presence of cGMP (Fig. 4F), reflecting a change in the conformation of the protein. On the other hand, neither the L267A nor the F295A mutant biosensor showed a discernable change in BRET in the presence of 1 µM cGMP (Fig. 4F). These results indicate that the mutations resulted in either a decrease in the affinity of the GAFa domain for cGMP or alteration in the structure of the domain in a way that cGMP binding does not induce a conformational change that could be detected using BRET. To determine which of these possibilities is responsible for the lack of an increase in BRET, we performed dose-response experiments with a range of cGMP concentrations. This revealed a cGMP concentration-dependent increase in BRET of WT GAFa domain biosensor, with an *EC*_50_ value of 19 ± 13 nM (Fig. 4G,H), which is similar to the previously reported value [49]. Interestingly, both the L267A and F295A mutant GAFa domain biosensors showed increase in BRET similar to that of the WT protein, albeit at much higher concentrations of cGMP with apparent *EC*_50_ values of ∼110 ± 74 and ∼194 ± 71 µM for the L267A and the F295A mutant, respectively (∼5800- and 10,000-fold higher for the L267A and the F295A mutant than the WT protein; with *p* = 10^−4^ for WT vs L267A and *p* < 10^−4^ for WT vs F295A; the actual *EC*_50_ values are likely to be higher as the BRET increases did not saturate at the highest concentration of cGMP tested here) (Fig. 4G,H). Together, these results suggest that the L267A and F295A mutations, although positioned away from the cGMP binding site, decreased the cGMP affinity of the GAFa domain.

Given that the L267A and F295A mutants showed a reduction in the basal BRET, we attempted to determine the impact of these mutations on the thermal stability of the GAFa domain. We posited that thermal denaturation of the GAFa domain would lead to a decrease in BRET of the biosensor constructs.[44] For this, we utilized the thermally stable fluorescent protein, mNeonGreen, as the BRET acceptor and the NLuc luciferase protein as the BRET donor, instead of GFP^2^ and RLuc due to their low thermal stability.[51] We generated plasmid expressing mNG-GAFa-NLuc biosensors, either WT or L267A or F295A mutant, and prepared lysates after transfecting HEK293T cells. Lysates were then incubated at a range of temperatures (ranging from 30 to 80 °C) for 10 min, following which BRET was measured. BRET data were fitted to a Boltzmann sigmoidal model to determine the *T*_m_, the temperature at which the relative BRET ratio is half of its maximum value (Supporting Figure 7). This revealed *T*_m_ values of 65.23 ± 0.55, 63.51 ± 0.13, and 59.93 ± 0.46 °C for the WT, L267A and F295A mutant GAFa domains (*p* = 0.0062 and 0.0002 for WT vs L267A and WT vs F295A, respectively) (Fig. 4I), suggesting that the mutations resulted in a minor, although statistically significant, decrease in the thermal stability of the GAFa domain. Importantly, the temperature-dependent decrease in BRET of the L267A was found to be steeper in comparison to the WT and F295A mutant GAFa domains (slope of −2.64 ± 0.25, −1.04 ± 0.10 and −2.88 ± 0.20 for WT, L267A and F295A mutant, respectively; *p* = 0.0005 for WT vs L267A mutant) (Fig. 4J). These results suggest that the L267A mutation leads to a resistance to thermal denaturation in the initial phases (below 60 °C), in a broad agreement with the decreased *R*g, and thus, a structural compactness, observed in MD simulations of the apo L267A mutant (Fig. 3D). Overall, these results suggest that lower basal BRET and higher cGMP apparent *EC*_50_ of cGMP-induced conformational change observed for the L267A and F295A mutant is likely not due to a decrease in the thermal stability of the proteins but instead likely reflects a change in their conformation and structural dynamics.

### L267A and F295A mutations induce a conformational change and significantly reduce the cGMP-binding induced conformational change in the full-length PDE5A2

Having elucidated the impact of the L267A and F295A mutations in the isolated GAFa domain, we aimed to investigate their effect on the cGMP-binding induced conformational change in the full-length PDE5 protein. For this, we incorporated the mutations in our previously described BRET-based, full-length PDE5A2 biosensor (Fig. 5A) that reports a conformational change upon cGMP binding to the GAFa, but not the catalytic, domain in the protein [39]. Western blot analysis revealed similar expression of the WT and mutant biosensor constructs (Fig. 5B; Supporting Figure 8). Bioluminescence spectral analysis revealed a second peak corresponding to GFP^2^ suggesting resonance energy transfer from RLuc in the case of the WT PDE5A2 biosensor (Fig. 5C). Importantly, as seen with the isolated GAFa domain biosensor, a decrease in the resonance energy transfer was observed for L267A and F295A mutant biosensors (Fig. 5C), leading to a significant decrease in basal BRET, as compared to the WT biosensor (Fig. 5D) (0.222 ± 0.009, 0.115 ± 0.002, and 0.122 ± 0.004 for WT, L267A, and F295A, respectively, *p* < 10^−4^ for both WT vs L267A and WT vs F295A), suggestive of a conformational change in the mutant proteins. Further, dose-response experiments with a range of cGMP concentrations revealed both a decrease in the maximum BRET change (−26.60 ± 3.42, −17.43 ± 1.31, and −12.60 ± 2.37% for WT, L267A, and F295A, respectively) and a right shift in the dose-response curve in the mutant biosensors (Fig. 5E), with *EC*_50_ values of 73 ± 20 nM, 181 ± 30 nM, and 3.4 ± 2.1 µM for WT, L267A, and F295A, respectively, *p* = 0.0071 and 0.0496 for WT vs L267A and WT vs F295A, respectively (Fig. 5F). These results indicate a change in the conformation in the absence of cGMP and a substantial decrease in cGMP affinity of the GAFa domain in the L267A and F295A mutant full-length PDE5A2 biosensors (Fig. 5F). Interestingly, the increase in the EC_50_ was greater in the F295A mutant compared to the L267A mutant indicating a more detrimental effect seen when mutating the F295 position in full-length PDE5A2.

**Fig. 5.**
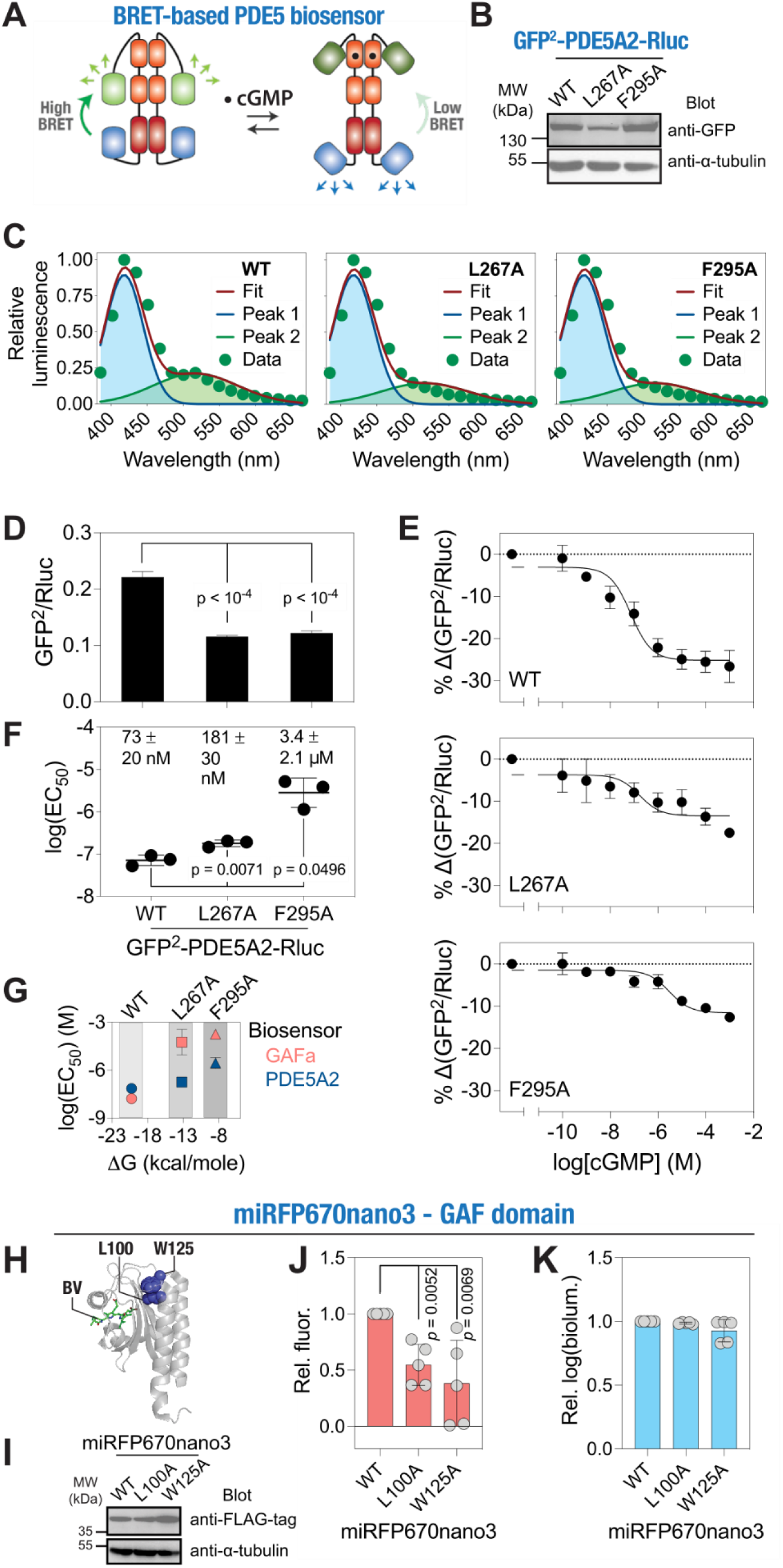
Mutation of distant, coevolving residues alters cGMP-induced allosteric regulation in the full-length PDE5. (A) Schematic showing the mechanism of the BRET^2^-based, full-length PDE5A2 conformational biosensor in the absence and presence of cGMP. (B) Western blot analysis of the cell lysates prepared from HEK293T cells transfected with either the WT or mutant full-length PDE5A2 biosensor plasmid constructs probed using an anti-GFP antibody showing the expression of biosensor constructs. (C) Graphs showing bioluminescence spectra obtained from lysates prepared from cells expressing the WT and mutant L267A and F295A mutant PDE5 biosensor constructs. Note the reduction in the GFP^2^ emission peaks in the L267A and F295A mutant PDE5 biosensors. Data shown are mean ± S.D. from a representative experiment, with experiments performed three times. (D) Bar graph showing the basal BRET values of the WT, L267A, and F295A mutant PDE5 biosensors. Note the significant reduction in the basal BRET values of the L267A and F295A mutants compared to the WT GAFa domain. (E) Graphs showing a percentage decrease in BRET values of the WT, L267A, and F295A mutant PDE5 biosensors after 30 min incubation with the indicated cGMP concentrations. Note the decrease in the maximum %change in BRET as well as the rightward shift in the cGMP dose-response curves of the mutant biosensors. (F) Graph showing log(EC_50_) values of cGMP-induced conformational change for the WT, L267A and F295A mutant GAFa domains in the full-length PDE5. Inset, values on top indicate the EC_50_ of cGMP-induced conformational change for the respective GAFa domains in the full-length PDE5 biosensor. For (D), (E), and (F), data shown are mean ± S.D. from three experiments, with each experiment performed in triplicates. (G) Graph showing log(EC_50_) (mean ± S.D.) values of cGMP-induced conformational change in the GAFa domain and full-length PDE5 proteins against ΔG (mean ± S.D.) values obtained from the MD simulation runs of GAFa domain (WT and mutants). (H) Cartoon representation of the miRFP670nano3 GAF domain (PDB: 7LSC) highlighting distant coevolving residues equivalent to PDE5 L267 and F295 (L100 and W125, respectively) and the ligand, biliverdin (BV). (I) Western blot analysis of the cell lysates prepared from HEK293T cells transfected with either the WT or mutant miRFP670nano3 plasmid constructs probed using an anti-FLAG antibody showing the expression of biosensor constructs. (J,K) Graphs showing relative fluorescence (J) and bioluminescence (K) of the WT, L100A and W125A mutant miRFP670nano3. Data shown are mean ± S.D. obtained from five independent experiments. *p*-values shown in panel (J) were obtained from Student’s t-test (unpaired, equal variance). For all tests, a *p*-value of <0.05 was considered statistically significant.

Having determined the *EC*_50_ values of cGMP-induced conformational change in both the isolated GAFa domain as well as the full-length PDE5A2, we compared these values with the free energy changes (*ΔG*) of cGMP binding to the WT and mutant GAFa domains determined from the MD simulation trajectories. This revealed a general decrease in the free energy changes of cGMP binding with both L267A and F295A mutant GAFa domain, although the effect was less pronounced in the case of the L267A mutant (Fig. 5G). Further, a clear difference could be observed in the impact of the two mutations in the isolated GAFa domain and the full-length PDE5A2 biosensors in terms of EC_50_ of cGMP-induced conformational change (Fig. 5G), suggesting that additional domains such as the GAFb and the catalytic domain in the full-length proteins (Fig. 5A) have a role in the cGMP-induced conformational change in the GAFa domain.

To further support the important role of the two coevolving residues in GAF domains, multiple sequence alignment of human cyclic nucleotide-binding PDEs revealed that these two amino acids are conserved in both GAFa and GAFb domains of these PDEs (Supporting Text 5). Additionally, analysis of variants at these positions in several PDE genes available in the Genome Aggregation Database (gnomAD) revealed a potentially harmful impact of the variations, further supporting their critical role in the functioning of the associated PDEs (Supporting Table 1) (See Supporting Methods and Supporting Results).

### Mutation of the distant coevolving residues in the fluorescent GAF domain protein, miRFP670nano3, decreases its fluorescence

Having established the impact of the distant coevolving residue positions 267 and 295 in the GAFa domain of PDE5, we aimed to determine the general applicability of these findings with respect to ligand binding in GAF domains in general. For this, we analyzed the structures of a variety of GAF domains available in the RSCB database and found the two residue positions to be in close proximity to each other and distant from the bound ligand, in cases where the structures contained a ligand, (Supporting Figure 9), suggesting a conserved role of the two residues. To confirm the role of the two coevolving residues in GAF domain function, we utilized a recently reported GAF domain fluorescent protein, miRFP60nano3.[52-54] It is a small, single-domain, engineered near-infrared fluorescent protein derived from cyanobacteriochrome photoreceptor GAF domain (Fig. 5H) that covalently binds the chromophore, biliverdin.[52-54]

We expressed the WT and L100A and W125A mutants (equivalent to the L267A and F295A GAFa domain mutations, respectively; based on structural alignment) (Fig. 5H), in HEK293T cells. We included a C-terminal fusion of a luciferase protein, picALuc(E50A)[41], for monitoring protein expression levels. Protein expression was also assessed using Western blot assay, which showed similar expression of the proteins (Fig. 5I; Supporting Figure 10). Fluorescence measurements revealed a significant reduction in cells expressing the L100A and W125A mutant miRFP670nano3, as compared to the WT protein (relative fluorescence of 1.00 ± 0.00, 0.55 ± 0.18, 0.38 ± 0.38 for WT, L100A, and W125A, *p* = 0.0052, 0.0069 for WT vs L100A and WT vs W125A, respectively) (Fig. 5J) while the expression of the proteins was found to be similar (Fig. 5I,K). These results indicate that the distant, coevolving residues are likely required for the miRFP670nano3 GAF domain to bind its ligand, biliverdin, required for its fluorescence activity and thus, confirm that the two coevolving residues are important for GAF domain function.

### Elucidating cGMP-induced allosteric activation of PDE5 GAFa domain

Having established the impact of mutations of the distant, coevolving residues in GAF domain function in the PDE5 GAFa domain and miRFP670nano3 proteins, we aimed to better understand the mechanism of cGMP-induced conformational change and the impact of the L267A and F295A mutations in the process through analysis of the MD simulation trajectories of the apo and holo GAFa, WT and mutant, domains. Specifically, we measured the distance between sheet β3 and helix α4 surrounding the cGMP binding site (Fig. 6A, left panel), as the two secondary structural elements act as a ‘lid’ that gates the cGMP binding site and have been reported to ‘close in’ upon cGMP binding (cGMP binding signal) [23]. Additionally, we measured the distance between helices α2 and α6 that are located at the N- and C-termini of the domain, respectively, and have been found to move closer upon cGMP binding, and thus, likely represents the allosteric activation signal (Fig. 6A, right panel). Following measurement of the two distances, i.e. cGMP binding associated (β3-α4 distance), and allosteric activation associated (α2-α6 distance), we pooled the distances from all three 1000 ns MD simulation trajectories and plotted the probability densities of the distances for both apo and the holo GAFa, WT and L267A and F295A mutant, domains (Fig. 6B). This analysis revealed that the β3-α4 and α2-α6 distances in the apo, WT GAFa domain largely centered around 11 and 13 Å, respectively, while the distances in the holo, WT GAFa domain centered around 10 and 9 Å, respectively, (Fig. 6B), indicating that cGMP binding to the WT GAFa domain results in a decrease in both the β3-α4 and α2-α6 distances, i.e. a ‘closing’ of the cGMP binding site ‘lid’ and a ‘closing in’ of the N- and C-terminal α-helices. The latter is likely the basis for the increase in BRET in the presence of cGMP that we observed with the WT GAFa domain biosensor (Fig. 4F,G). Similar findings were reported for the isolated GAF domain of Anabaena adenylyl cyclase utilizing amide hydrogen/deuterium exchange mass spectrometry [7].

**Fig. 6.**
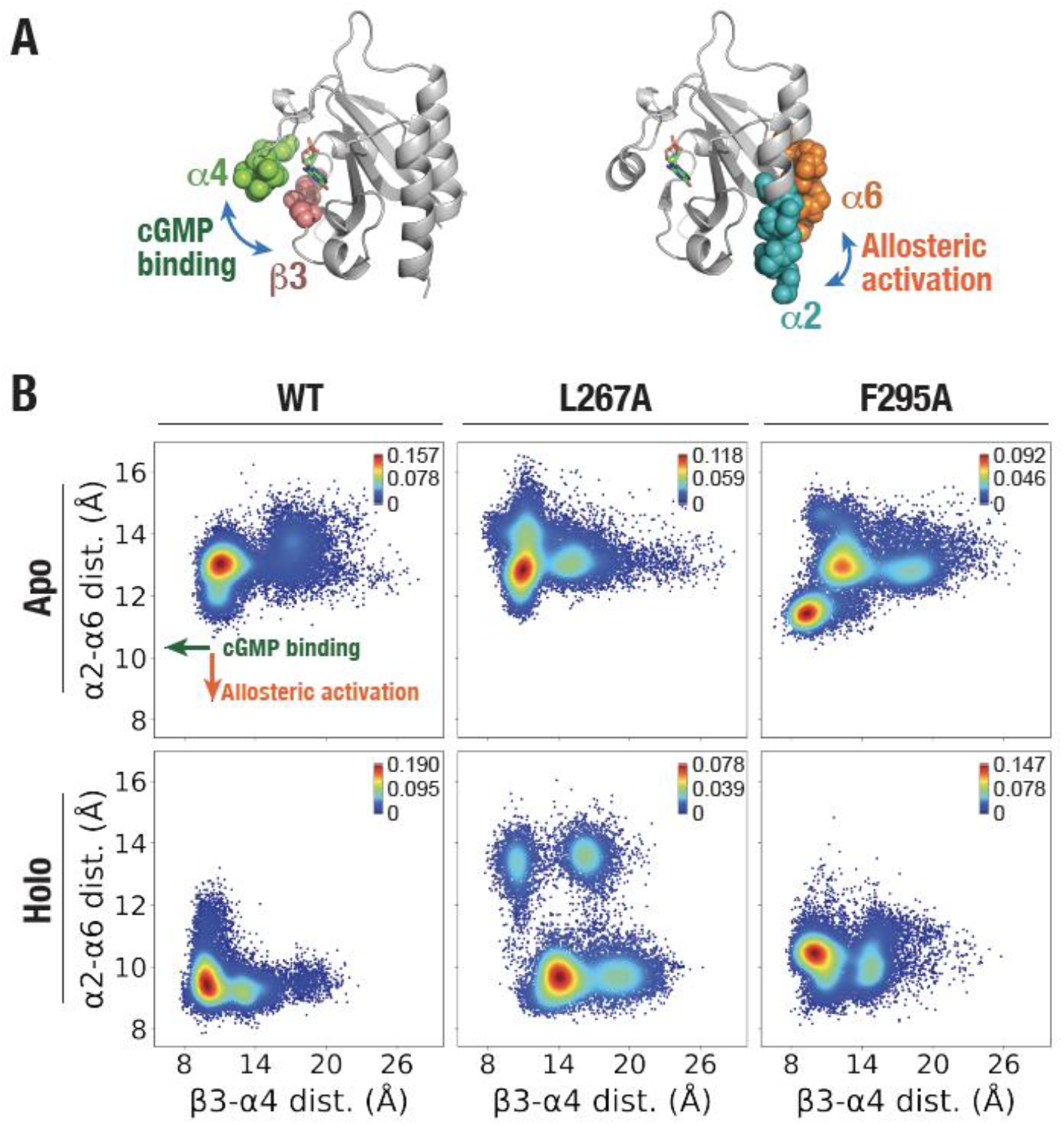
Altered ligand binding site and allosteric activation upon mutation of distant, coevolving residues in the PDE5 GAFa domain. (A) Cartoon representation of the GAFa domain highlighting helix α4 and sheet β3 from the ligand binding site that ‘closes’ upon cGMP binding (left panel) and N-terminal helix α2 and C-terminal helix α6 that move closer upon cGMP binding (right panel). (B) Graphs showing probability distribution (kernel density estimation) of distances between helix α4 and sheet β3 (x-axis) and N-terminal helix α2 and C-terminal helix α6 (y-axis) of the apo (upper panel) and holo (lower panel) GAFa domain (WT and mutants) determined from a cumulative of three independent, 1000 ns long MD simulation trajectories. Color bars, probability densities.

In the apo L267A mutant GAFa domain, both the β3-α4 and α2-α6 distances were centered similar to that of the WT GAFa domain (Fig. 6B). In the holo L267A mutant, the α2-α6 distance was decreased (from 13 to 9.5 Å) to levels similar to that of the WT GAFa domain (9 Å). However, the β3-α4 distance was found to increase (from 11.5 to 14 Å), instead of decrease as seen with the WT GAFa domain (Fig. 6B). This suggests that the L267A mutation resulted in the opening of the ‘lid’ of the cGMP binding site and thus, likely, enhancing the unbinding of cGMP from the protein leading to an increase in the *EC*_50_ of cGMP-binding.

On the other hand, in the apo F295A mutant GAFa domain, two separate density centers (one with β3-α4 and α2-α6 distances of 10 and 11.5 Å, respectively, and another with distances of 13.4 and 13 Å, respectively) were observed (Fig. 6B), suggesting a destabilization effect of the mutation in the apo state affecting both the lid’ of the cGMP binding site and the terminal helices. However, in the holo state, in which the WT GAFa domain assumed a more compact form in comparison to the apo form, the F295A mutant displayed a density center that is similar to the WT holo state with regards to the β3-α4 distance (10 and 10 Å for the F295A and WT GAF domains, respectively), but the α2-α6 distance found to be similar to that of the major center seen in the apo state of the protein (11.5 and 11.5 Å, respectively) with a loss of the second center observed in the apo state, which was higher than the distance observed with the holo WT GAFa domain (11.5 and 9.5 Å, respectively) (Fig. 6B). These results suggest that the cGMP binding may not lead to the effective transduction of the allosteric signal, which agrees with an increase in the *EC*_50_ of cGMP-induced conformational change determined using the GAFa biosensor (Fig. 4F,G). Overall, these results illustrate the impact of L267A and F295A mutations in the GAFa domain conformational landscape and point to a critical role played by the two residues in cGMP-induced conformational change in the GAFa domain, and likely impact this through two distinct mechanisms.

## Discussion

GAF domains are among the most conserved molecular switches that are abundantly present in different proteins across a diverse range of species. In this study, we used SCA to identify a cluster of coevolving residue positions in GAF domains. The SCA method has been successfully applied in a number of cases, including decoding the functional evolution of enzymes [55] and their metal binding specificities [56], discrimination between correct and incorrect protein folds that are based on de-novo structure prediction [57], and predicting allosteric networks in proteins [58-62] as well as functional RNA molecules [63]. We identified two residue positions that showed the highest coevolutionary score and are located distant from the ligand binding site. Mapping the two residue positions on GAF domain structures from different species showed that the two residues are juxtaposed and may play a role in the allosteric function of the domain. We used the GAFa domain of PDE5 to investigate the role of the two residue positions (L267 and F295) in the domain. Our in-silico analysis of the apo and holo forms of the domain, as well as in vitro assays shed some light on the functional role of the two positions on the ligand-binding induced conformational change in the domain. Mutating the L276 residue, which is located in the sheet β5, induces a structural change in the domain that we detected experimentally as a decrease in BRET. This change in the conformation is associated with a decrease in the structural dynamics of the domain and converts the apo state into a more compact form, as can be inferred from the RMSD, RMSF, and *R*g analysis of the MD trajectories. This, in turn, could potentially affect the ligand binding as structural flexibility of the apo form is an important feature that facilitates ligand binding. To further support the idea of these structural alterations, our in vitro assays showed that the L267A mutant is more resistant to the thermal unfolding with increasing temperature, although the *T*_m_ value was slightly decreased compared to the WT. Additionally, our MD simulation analysis showed that, in the holo form, the L267A mutant results in the opening of the α3-β4 “lid” that gate the cGMP binding site, making it difficult to retain the ligand in the binding site. This was reflected by a decrease in the ligand SASA, interaction energy, hydrogen bond formation, and free binding energy in the L267A mutant domain compared to the WT. These computational findings were further confirmed experimentally using an L267A mutant BRET-based GAFa domain biosensor, which showed more than 5000-fold increase in the EC_50_ of the cGMP-binding induced conformational change, compared to the WT GAFa domain.

On the other hand, the F295 residue is located on the β6/α5 loop near the terminal helix (α5) that transmits the allosteric signal to the catalytic domain of the protein. Our results show that the F295A mutation induces a structural change in the domain that was detected experimentally as a decrease in the basal BRET ratio of the F295A mutant GAFa-domain biosensor. Based on the MD simulation trajectory analysis, this mutation results in the disruption of the inter-residue interactions that are originally formed between this position and neighboring residues, including the L267 residue, in the WT GAFa domain. This results in an increase in the structural dynamics of the domain, as can be inferred from the RMSD analysis, and, in the holo form, in a less compact structure as indicated by *R*g analysis. Specifically, the F295A mutation increases the α2-α6 distance in the holo form, suggesting that it interferes with the allosteric signal transduction to the catalytic domain following ligand binding. These results were further confirmed experimentally using an F295A mutant BRET-based GAFa-domain biosensor, which showed more than 10000-fold increase in the EC_50_ of the cGMP-induced conformational change as compared to the WT GAFa domain.

Overall, these results indicate that the two mutations negatively impact the cGMP-binding -induced conformation change in the isolated GAFa domain. The L267A mutation appears to directly affect ligand binding through a decrease in the flexibility and an increase in the rigidity of the apo form – both of which are important for ligand binding in the apo form - as well as by opening of the α3-β4 “lid”, that gate the cGMP binding site, in the holo form making it difficult to retain the ligand inside the binding pocket. On the other hand, the F295A mutation appears to negatively affect cGMP-binding induced conformational change by two mechanisms, both as a result of disrupting the local inter-residue hydrophobic interaction. First, this leads to the destabilization of the ligand inside the binding pocket, which directly and negatively affects ligand binding. Second, this disruption of the hydrophobic interaction resulting from mutation of this position leads to a change in the cGMP allosteric signal transmission to the terminal α5 helix, which extends to connect GAFa to GAFb domain in PDE5.

Furthermore, the structural changes induced by the two mutations in the isolated GAF domain were also recapitulated experimentally in the full-length enzyme as a decrease in the basal BRET ratio of the mutant PDE5-based biosensors. Moreover, the mutant biosensors displayed a suboptimal conformational change as indicated by the lower values of the maximal change in BRET ratio at the highest cGMP concentration tested. Additionally, there was a significant decrease in the ligand-binding induced conformational change in the mutant proteins as indicated by the cGMP EC_50_ values. This indicates the two positions play an important role in the allosteric function of the GAFa domain of PDE5. Importantly, there was a greater decrease in the EC_50_ in the case of the F295A full-length PDE5 mutant, which further supports the results reported here that the F295A mutant may also affect the allosteric signal transduction from GAFa domain to catalytic domain that follows ligand binding. Furthermore, we show that the functional role of the two positions is not restricted to the GAFa domain of PDE5 but extends to other GAF domains. In this regard, mutating the two positions in the engineered GAF domain fluorescent protein, miRFP670nano3, resulted in a decline in the fluorescent emission of the protein, likely due to a decrease in biliverdin binding.

In this regard, the near full-length structure of homodimeric PDE2A, which includes both the regulatory and catalytic domains, provided some insights into GAF domain-mediated regulation of PDE catalytic domain [64]. In the absence of cGMP binding, the two catalytic domains in the dimer occlude the substrate binding site in each other by means of a loop region (H-loop). Binding of the allosteric ligand, cGMP, to the GAFb domain causes the H-loop to ‘swing’ out of the catalytic pocket, allowing access to the catalytic site for the substrate, cGMP, molecules [64]. Relevant to this, a previous structural study of PDE6 tandem GAF domains (GAFa and GAFb), which shows the highest sequence and structural homology with tandem GAF of PDE5, reported cGMP-dependent structural changes in the GAFa domain that are similar to the GAFa of PDE5, including movement of the β1/β2 loop, the α2/α3 region, and α4 helix [65]. However, the same study did not report a cGMP-dependent change in the conformation and distance between the terminal helices, α2 and α6, which is the basis for the design of our in-house PDE5 GAFa-domain biosensor [38]. Additionally, the study suggested that allosteric signal transmission from the GAFa domain to the GAFb domain takes place at several sites where the two domains meet. This includes interactions from the GAFa β1/β2 loop to the GAFb β4/β5 and β6/α5 loops, as well as from the GAFa α2/3 region to the GAFb β6/α5 loop [65]. In fact, some of the reported PDE6 disease-causing mutations are located in the vicinity of the β1/β2 loop of GAFa [66-69], suggesting a disruption in the allosteric communication caused by these mutations in PDE6. On the other hand, our computational and biophysical investigations of the GAF domain provide evidence of additional, coevolving amino acid residues that are involved in the cGMP-induced conformational change. These coevolving residues are not only located distant from the cGMP binding site but also do not show interfacial interaction with the GAFb domain, suggesting a more complex and multimodal allosteric signal relay across cyclic nucleotide-binding PDEs than what has been proposed in previous studies.

Moreover, our results show that the two coevolving residues are invariant across the GAFa and GAFb domains of all human cyclic-nucleotide binding PDEs, and mutations at these two positions in human PDE variants that are reported in the gnomAD database were predicted to be potentially harmful to the associated PDE genes, strongly suggestive of a critical role played by these two residues in the GAF domain function, i.e. ligand binding and induction of allostery in the PDEs. On the other hand, the F295 position is a part of the NKFDE motif (N286, K287, F295, D299, E230, numbering according to PDE5A1) that distinguishes cyclic nucleotide-binding GAF domains from other GAF domains. Since they are conserved in cyclic nucleotide binding GAFs, the residues making up this motif were initially thought to directly interact with cyclic nucleotides [10, 70-72]. However, NMR and crystallographic structures later showed that residues making up this motif are not in direct contact with the cyclic nucleotide ligand, although this motif was still deemed necessary for stabilizing the GAF domain fold, cyclic nucleotide binding, and transmission of the allosteric signal to the catalytic domain [2, 11, 16, 70]. In fact, mutating the residues of the NKFDE motif into alanine, each individually, in the PDE2 GAFb domain abolished cGMP binding [46]. Similar observation was made upon mutation of the D residue into alanine in the GAFa domain of PDE11A4 [73]. Moreover, individually mutating N, K, and D residues of the NKFDE motif in PDE5 GAFa domain greatly decreased the cGMP binding affinity [70, 74]. With that being said, the role of the conserved NKFDE motif in cyclic nucleotide binding GAF domains remained enigmatic. It is worth noting that F295 is the only residue amongst the cluster of nine highly coevolving residue positions identified from our coevolutionary analysis (SCA) that is a part of the NKFDE motif, which critically highlights the significance of our study in understanding the ligand binding and allosteric regulation in GAF domain proteins. Overall, results presented here provide insights into the mechanism underlying allosteric regulation of GAF domains and can potentially pave the way for designing novel chemical modulators of GAF domain allostery.

## Conclusion

To conclude, we successfully utilized SCA to identify coevolving residue positions in GAF domains, two of which are distant from the ligand binding site but found to play a critical role in ligand-binding induced conformational allostery. Our analysis suggests the two residues could affect the GAF domain-mediated allostery by different mechanisms. The L267 position appears to be more involved in controlling ligand traffic to (apo) and from (holo) the ligand binding site in the domain. On the other hand, the F295 position is important for maintaining the local inter-residue interaction, and when disrupted it likely affects both ligand binding (holo) as well as signal transduction to the catalytic domain that follows ligand binding to the GAF domain. As such, our study proposes: first, a ligand-binding model in GAF domains, in which ligand access to/from the binding site is controlled by β3-α4 ‘lid’ that gate the ligand binding site, and therefore any factors that affect the stability/dynamics of these secondary structures are likely to affect the cGMP binding affinity, and second, a ligand-binding induced signal transduction model, in which the conformational change signal that follow ligand binding is transmitted to the catalytic domain through the C-terminal helix (α5) in the domain. Importantly, our data suggest that factors disrupting signal transduction in the domain could have serious detrimental effects on the allostery that are comparable, if not more, to factors that directly reduce ligand binding.

## Acknowledgements

This work is supported by a grant from HBKU Thematic Research Grant Program (VPR-TG01-007) and internal funding from the College of Health & Life Sciences, Hamad Bin Khalifa University, a member of the Qatar Foundation. W.S.A. is supported by scholarship from CHLS, HBKU. A.T. was supported by a Summer Research Fellowship Program from the Indian Academy of Sciences. A.M.G and T.A. are supported by a postdoctoral fellowship from CHLS, HBKU. Some of the computational research work reported in the manuscript was performed using high-performance computer resources and services provided by the Research Computing group in Texas A&M University at Qatar. Research Computing is funded by the Qatar Foundation for Education, Science and Community Development (http://www.qf.org.qa).

## Supporting Information

### Supporting Methods

#### Bioinformatic analysis

We performed multiple sequence alignment of GAF domain-containing human PDEs (PDE2, 5, 6, 10, and 11) to see if the two residues are conserved in other human GAF domains. Additionally, we utilized the Genome Aggregation Database (gnomAD) [75] to identify missense, insertion, or deletion variants that carry mutations at the selected coevolving residue positions. Pathogenesis potential of these variants was assessed using in silico prediction tools, including CAAD [76], REVEL [77], and PolyPhen [78]. We applied a cutoff of 20 [76, 79], 0.5 [80], and 0.908 [80] for the CAAD, REVEL, and PolyPhen prediction scores, respectively, to distinguish potentially harmful variants from benign ones.

The effect of mutating the coevolving residue positions on the protein stability was investigated by calculating Gibbs free energy changes (ΔΔ*G*) using in silico structure- and sequence-based prediction tools including DynaMut [81], mCSM [82], SDM [83], DUET [84], DeepDDG [85], iDeepDDG [85], Maestro [86], PoPMuSiC [87], I-Mutant 2.0 [88], CUPSAT [89], Mupro [90], and ENCoM [91]. To probe the effect of the mutations on the protein flexibility, the elastic network contact model (ENCoM)-based vibrational entropy difference (ΔΔSvib) between WT and mutants was calculated [91]. Solvent accessibility of the mutated residues was predicted using PoPMuSiC online tool [87]. Arpeggio webserver [92] was used to calculate and compare the number of intramolecular interactions formed by the coevolving residue positions. Moreover, the impact of the mutants on the domain function was predicted using the SNAP2 tool [93]. SNAP2 prediction scores reflect the likelihood of the specific mutation to alter the native protein function and range from −100 (strong neutral prediction) to +100 (strong effect prediction). The scores can be interpreted as strong signal of effect (>+50), weak signal (−50<score<+50), or strong signal of neutral or no effect (<-50). We also predicted the impact of the L267A and F295A mutation using the ESM variant analysis[94], which revealed a moderately disruptive effect for the L267A mutation and a highly disruptive effect of the F295A mutation on the PDE5 GAFa domain function.

### Supporting Results

#### Bioinformatics/genomic database analysis

Because our coevolutionary analysis suggested that the two coevolving residue positions may play a role in the GAF domain function, we wanted to see if these two residues are conserved in other human GAF domains. To achieve this, we performed multiple sequence alignment of GAF domain-containing human PDEs (PDE2, 5, 6, 10, and 11). The alignment showed that these two residues are indeed conserved across both the GAFa and GAFb domains of the cyclic-nucleotide binding human PDEs (Supporting Text 3), suggesting an essential role that these two residues may play in the function of the associated PDEs. To further confirm their importance, we assessed if there are any reported variants that involve mutations at these two locations in the PDE5 GAF domains. To achieve this, we searched for missense, insertion, or deletion variants in the gnomAD database [75]. We listed these variants in Supporting Table 1. Assessment of the deleterious effect of these variants using the in silico prediction tools CADD [76], REVEL [77], and PolyPhen [78] suggested that all variants have deleterious effects on the associated PDE genes, confirming the potentially essential role of these two residue positions on the associated PDE genes (Supporting Table 1).

Moreover, we utilized silico prediction tools to explore the impact of these two positions on the PDE5 GAFa domain stability, flexibility, and functionality. Individual mutation at either of the two positions was predicted by the SNAP2 tool to have a functionally deleterious effect. More specifically, our analysis showed that mutating either of the two residues results in a destabilizing effect on the domain. Additionally, there was an increase in the overall flexibility of the domain upon mutating the residue positions, with more flexibility observed for the F295A mutant (Supporting Table 2). This increase in flexibility was most observed for the α4 and terminal helices (Supporting Figure 11A). To explore how these effects are brought into display, we compared the number of contact interactions formed by these two positions in the WT and mutant domains. We found that mutating these two positions drastically affects the number of hydrophobic contacts they form with neighboring residues (Supporting Table 3; Supporting Figure 11B). Moreover, both residues are hydrophobic and, therefore, can be affected negatively by solvent accessibility. The PoPMuSiC tool predicted the 295 positions to have more solvent accessibility, compared to 0.0% accessibility for the 267 positions, which may contribute to the more profound effect observed upon mutating this position on the domain’s structure and function. Overall, our in-silico analysis shows that mutating the two coevolving residue positions is predicted to have drastic effects on the stability, flexibility, and functionality of the GAF domain facilitated by the excessive loss of hydrophobic contacts formed by the two residue positions.

### Supporting Texts

**Supporting Text 1:**
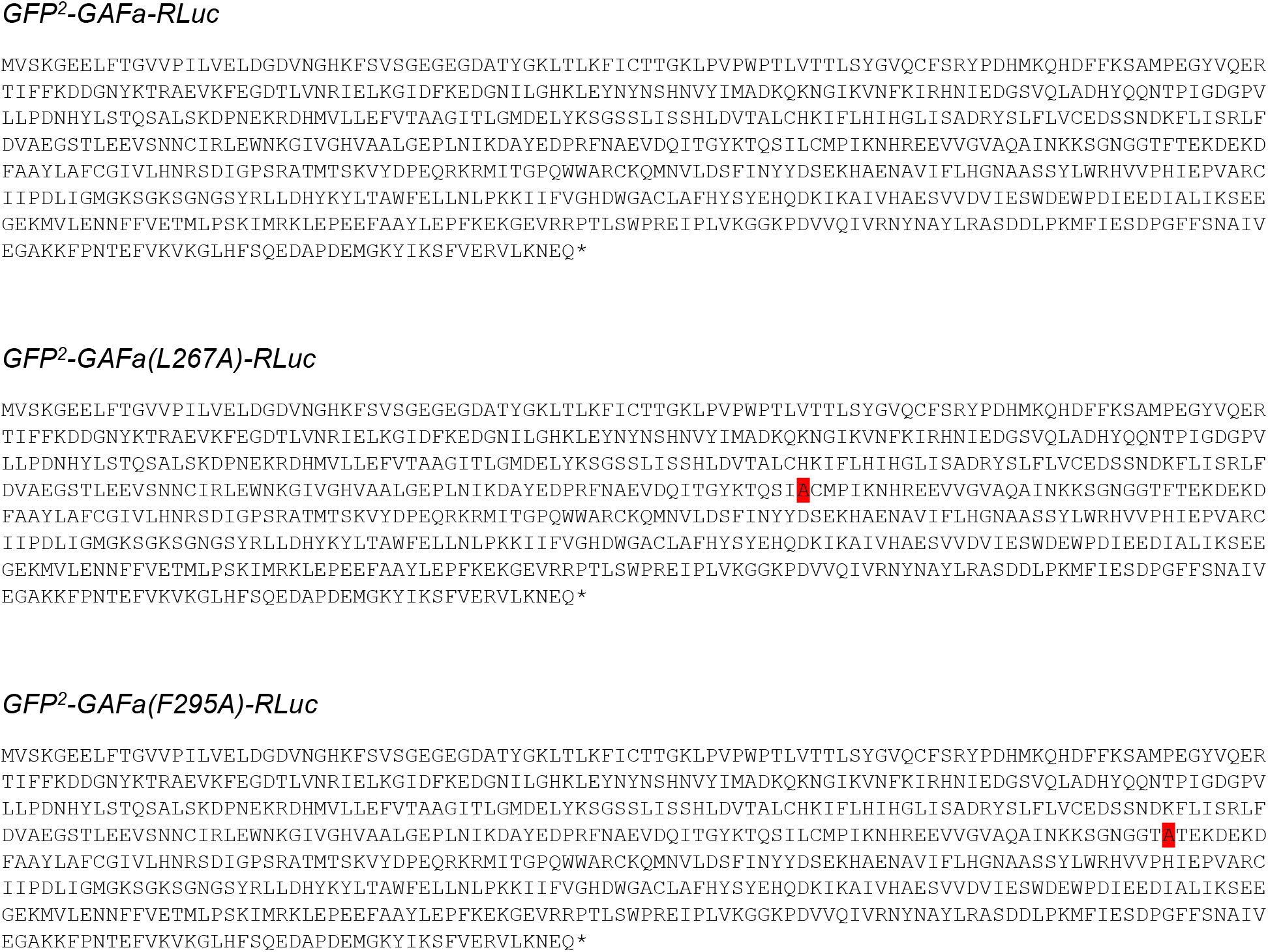
Protein sequence of GFP^2^-GAFa-RLuc constructs (mutations highlighted in red)

**Supporting Text 2:**
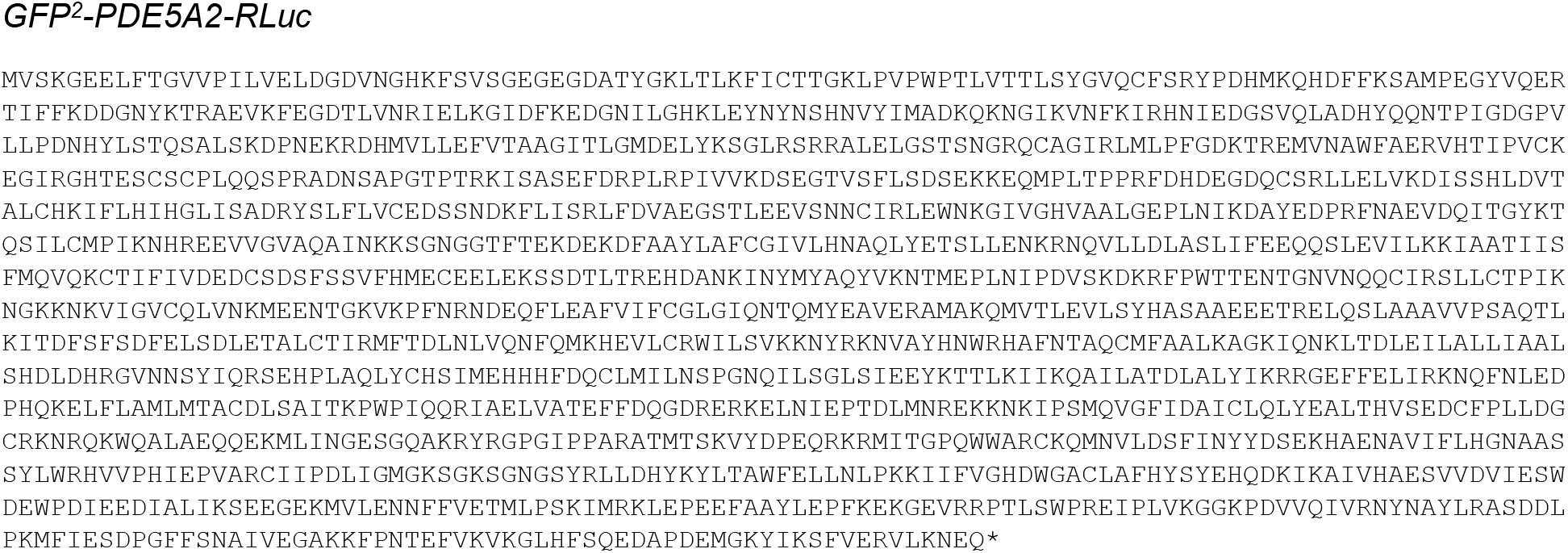
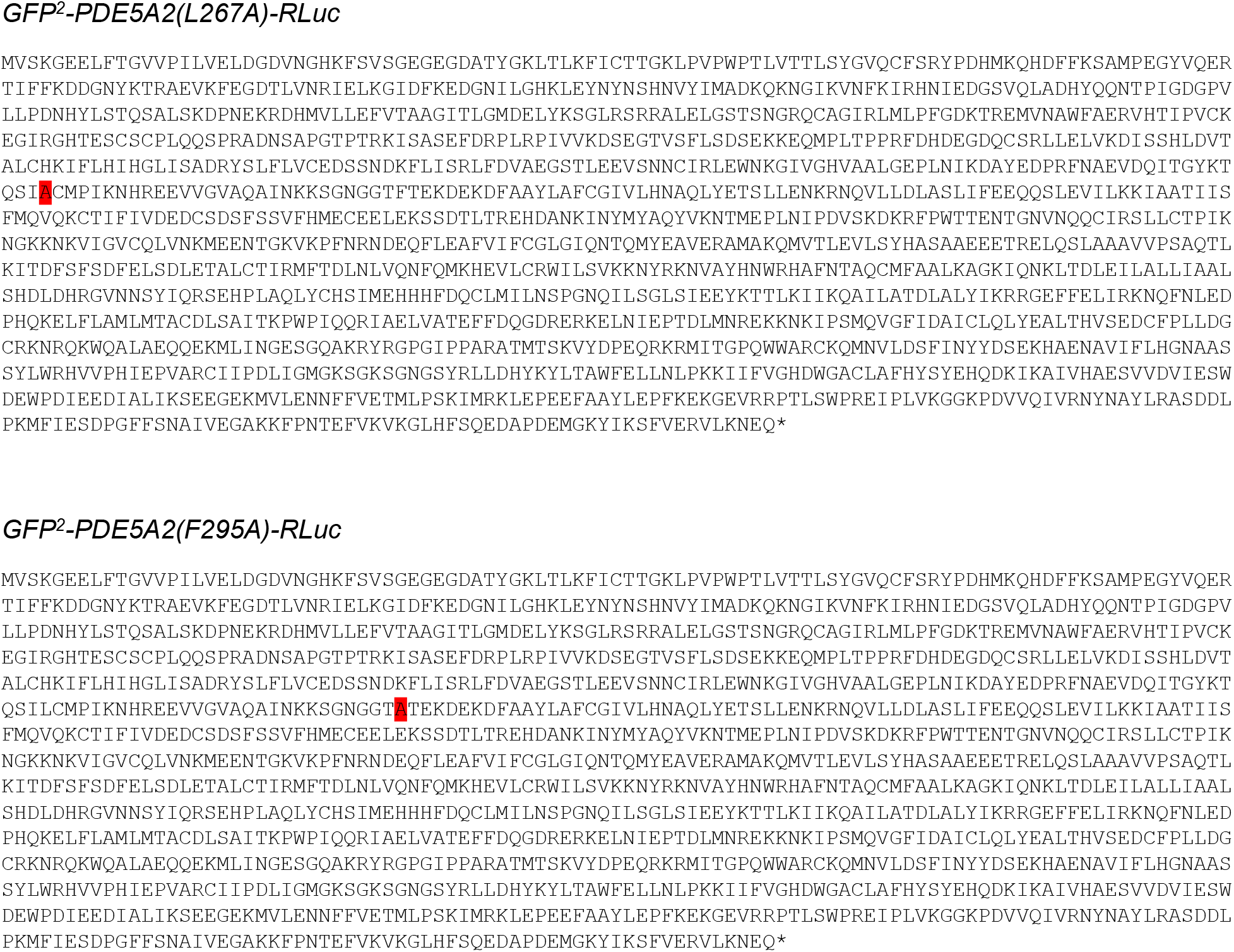
Protein sequence of GFP2-PDE5A2-RLuc constructs (mutations highlighted in red)

**Supporting Text 3:**
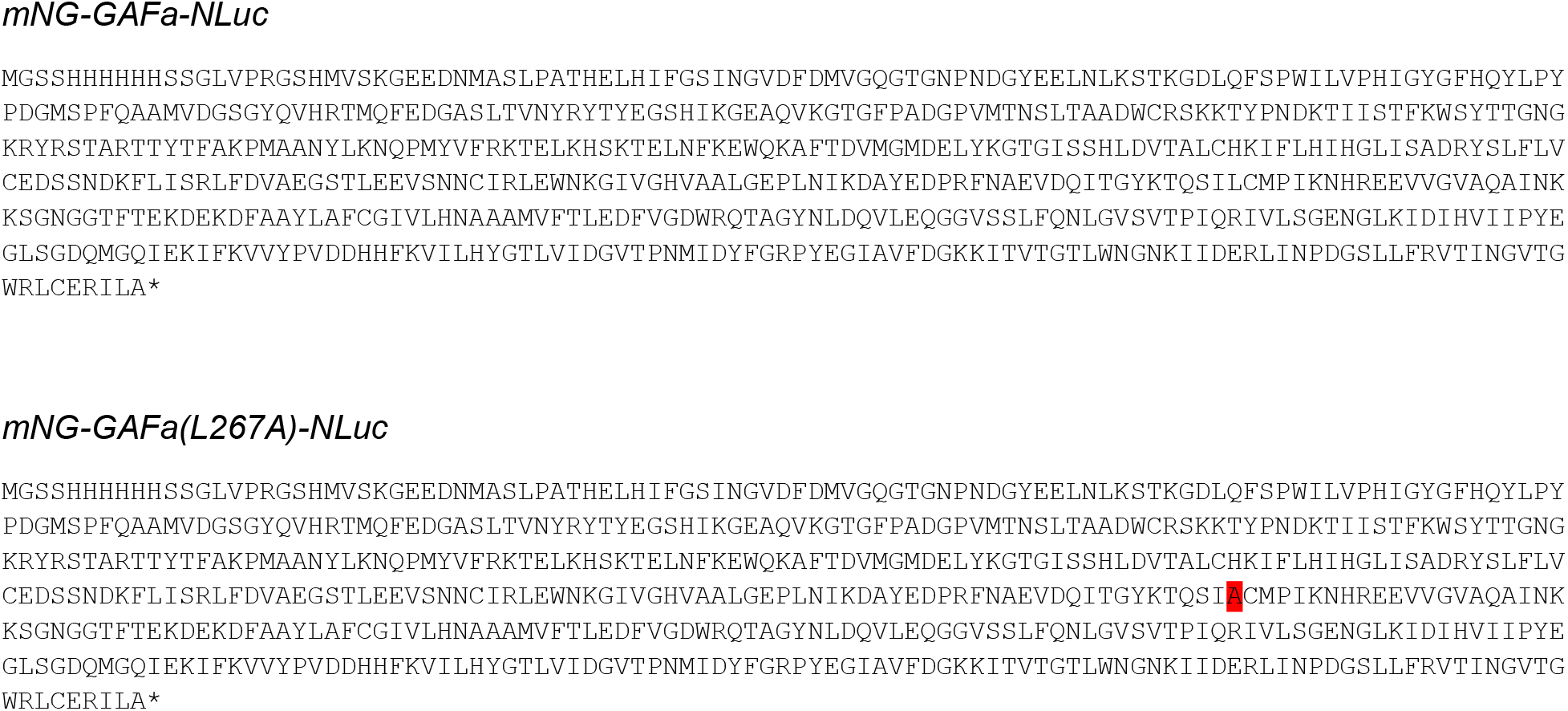
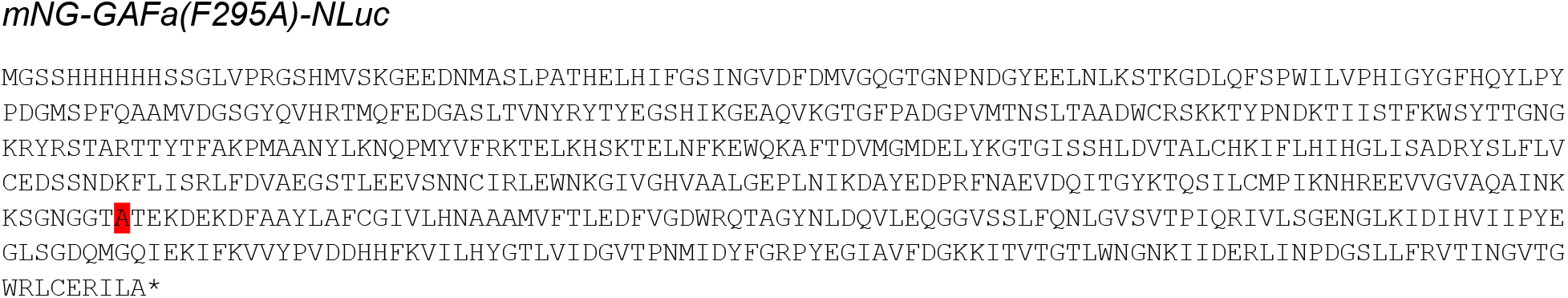
Protein sequence of mNG-GAFa-NLuc constructs (mutations highlighted in red)

**Supporting Text 4:**
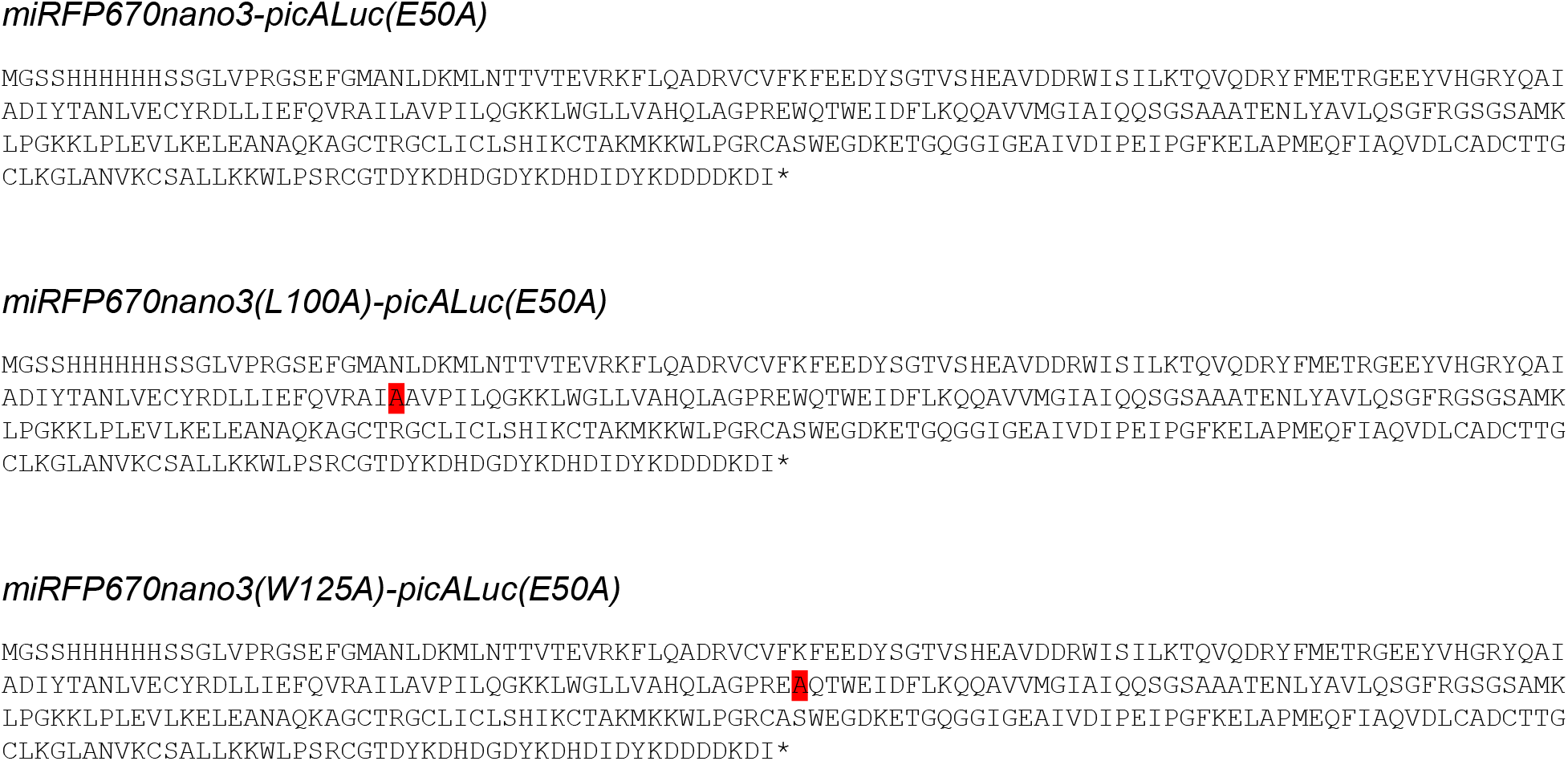
Protein sequence of miRFP670nano3 -picALuc(E50A) constructs (mutations highlighted in red)

**Supporting Text 5:**
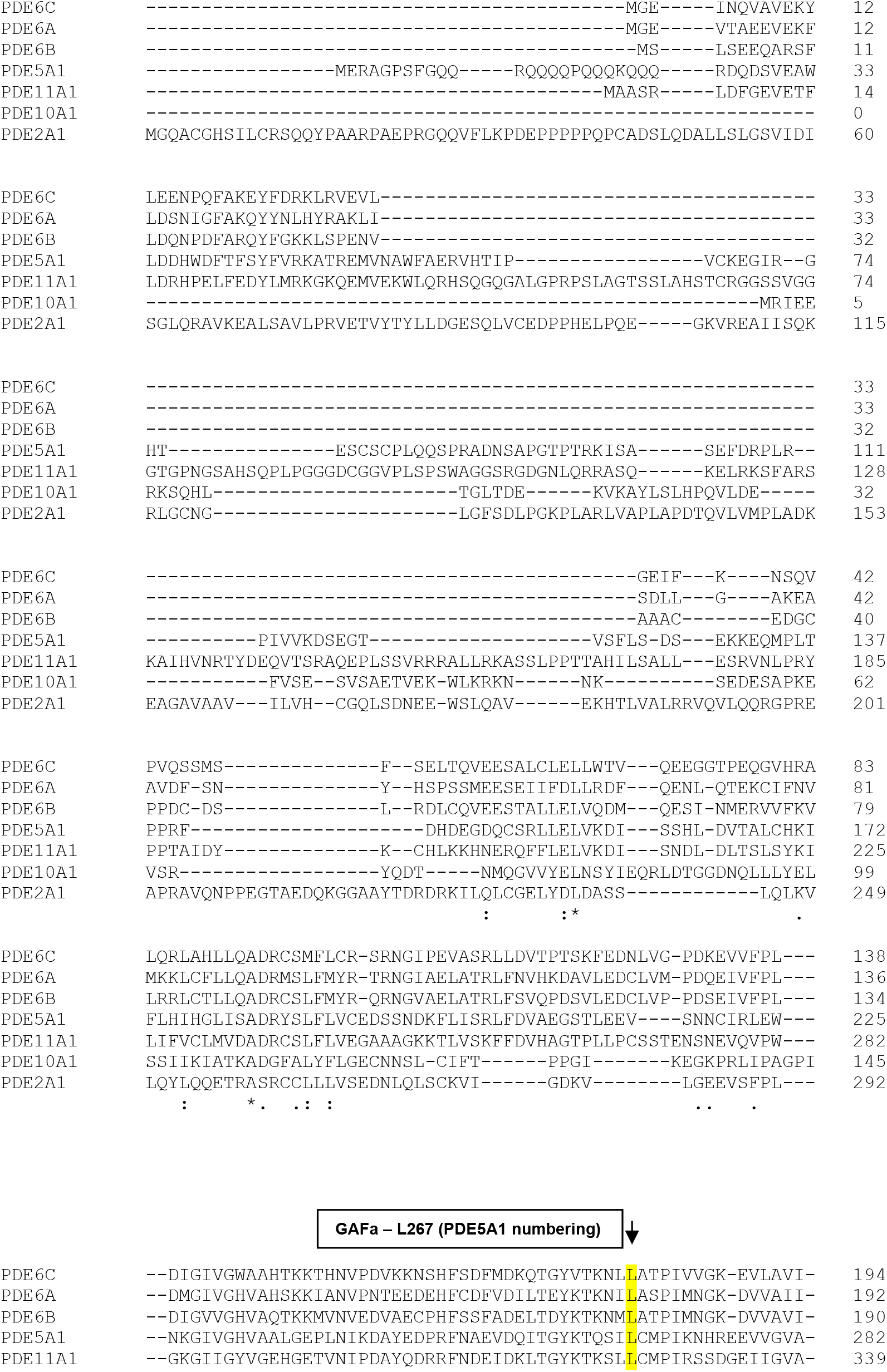
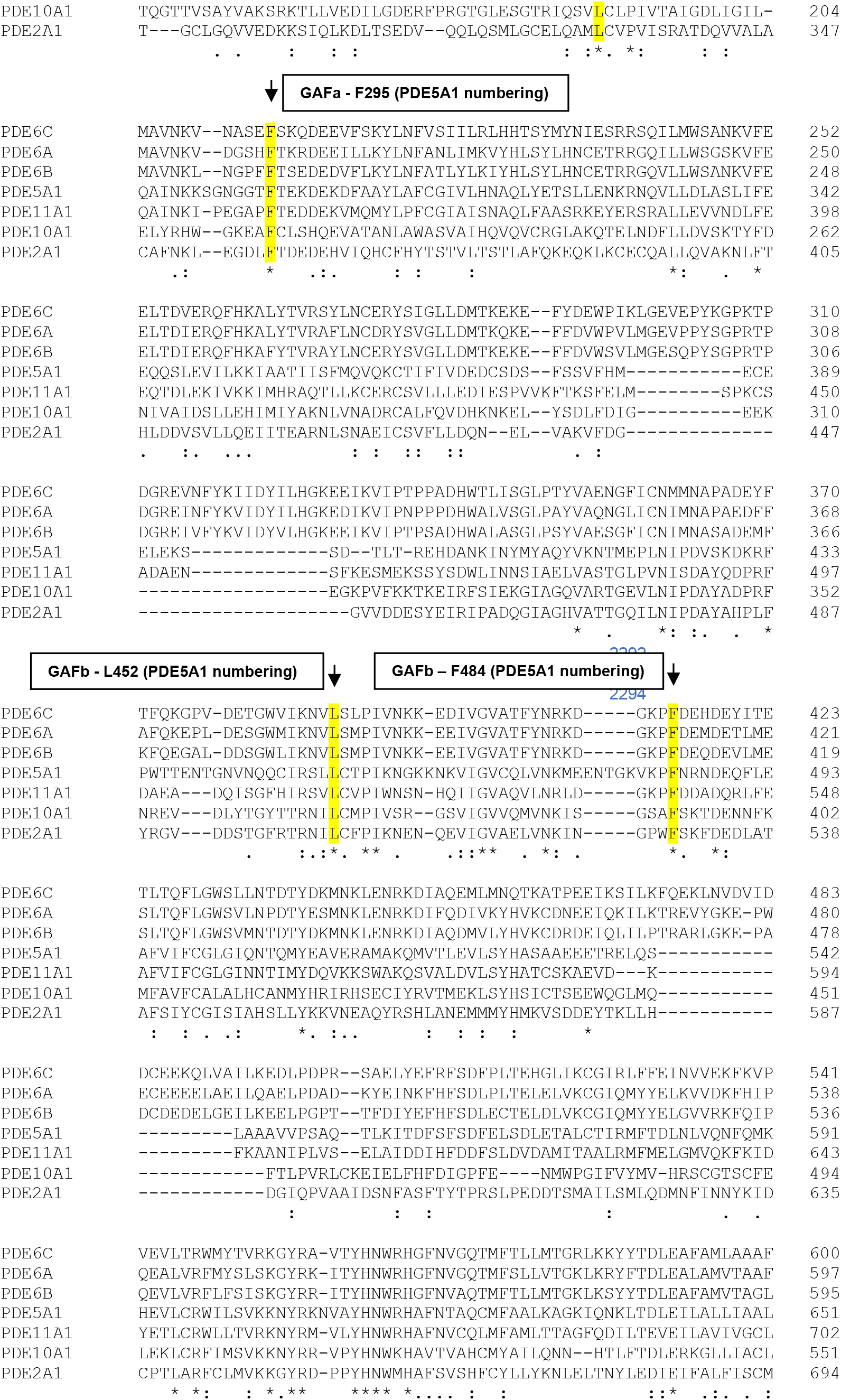
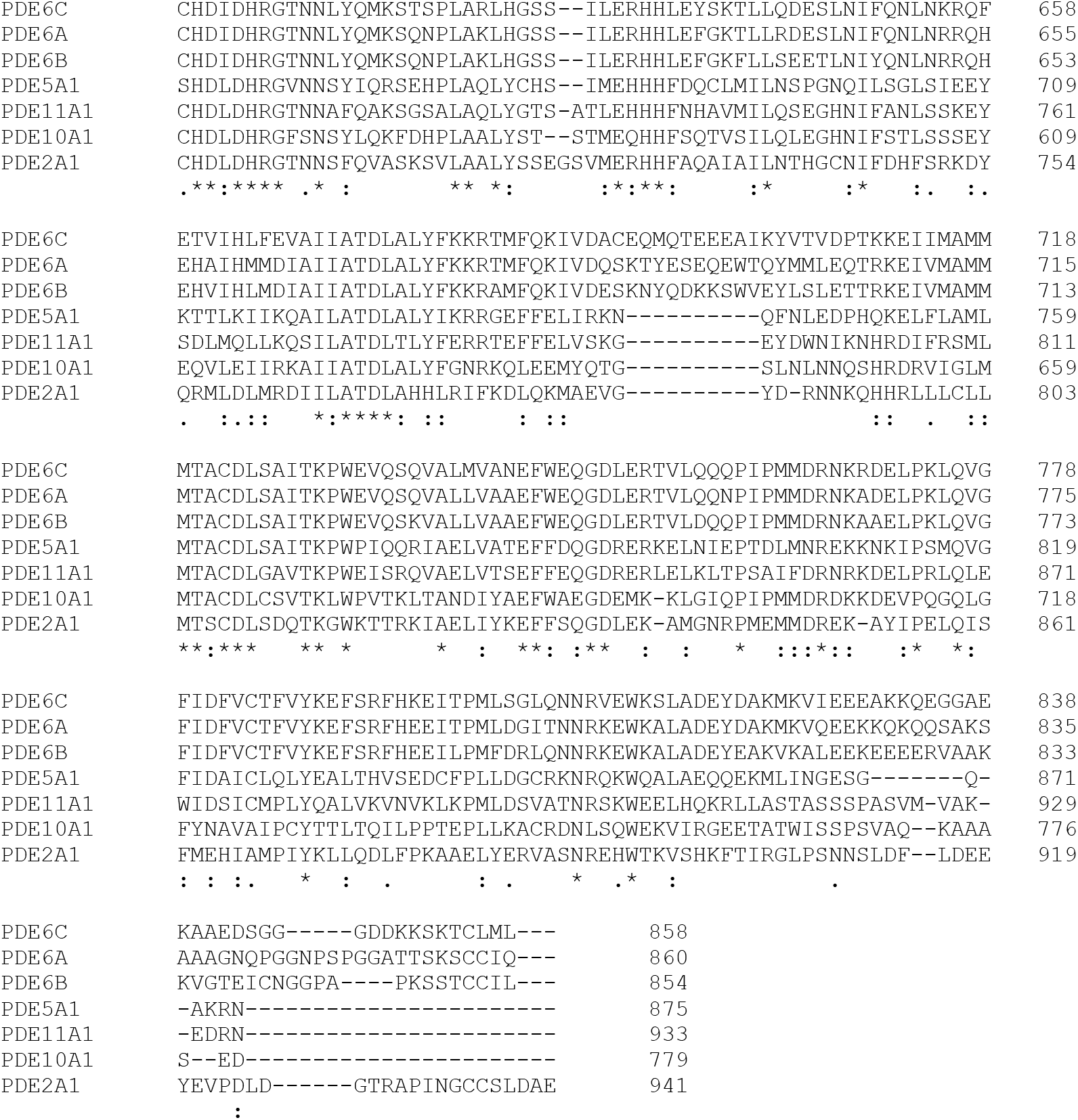
Multiple sequence alignment of GAF domain-containing human PDEs showing that both L267 and F295 positions are conserved in both GAFa and GAFb domains of all GAF domain-containing human PDEs.

**Supporting Text 6:**
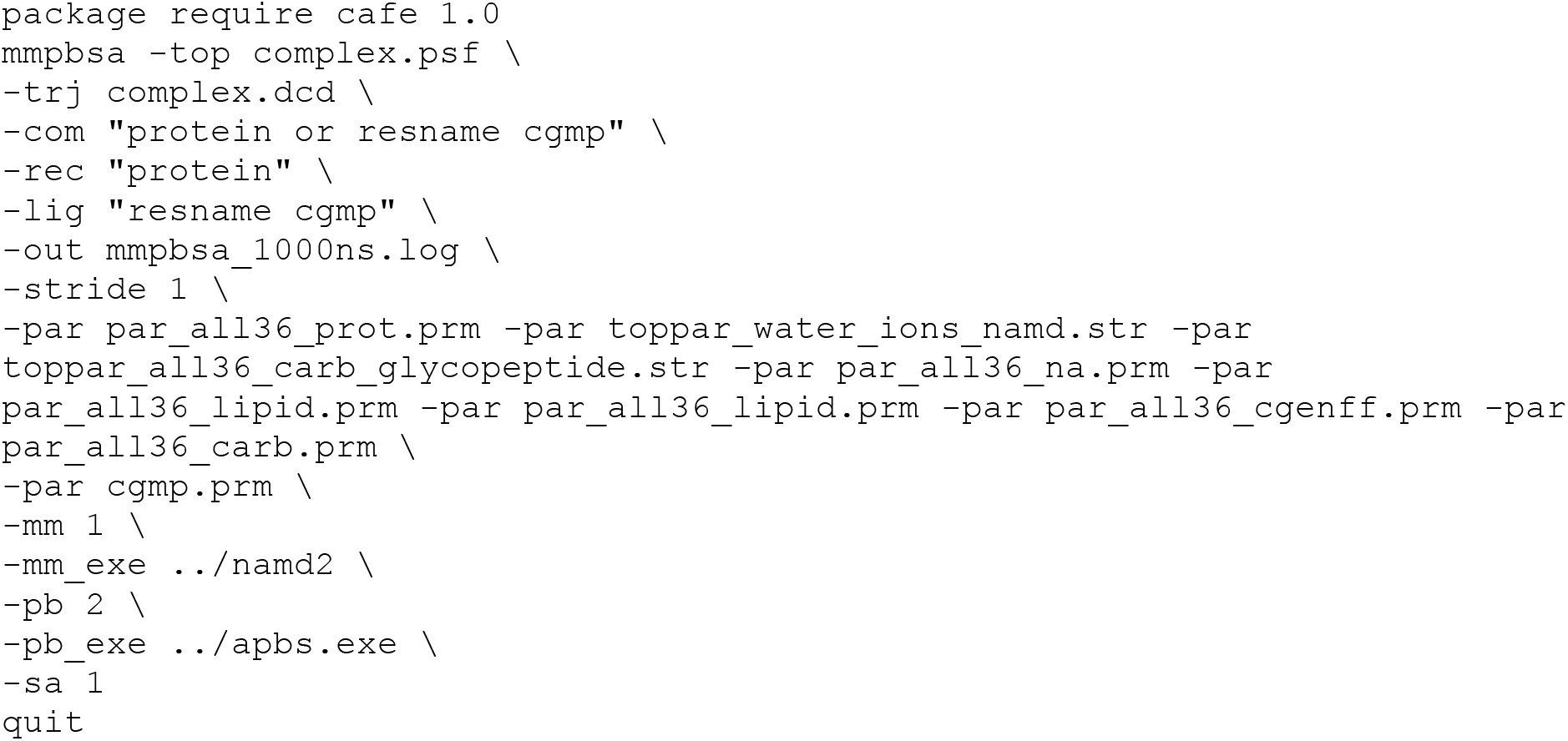
Tcl script used to calculate the free binding energy using the CaFE pluginin in VMD.

### Supporting Tables

**Supporting Table 1.**
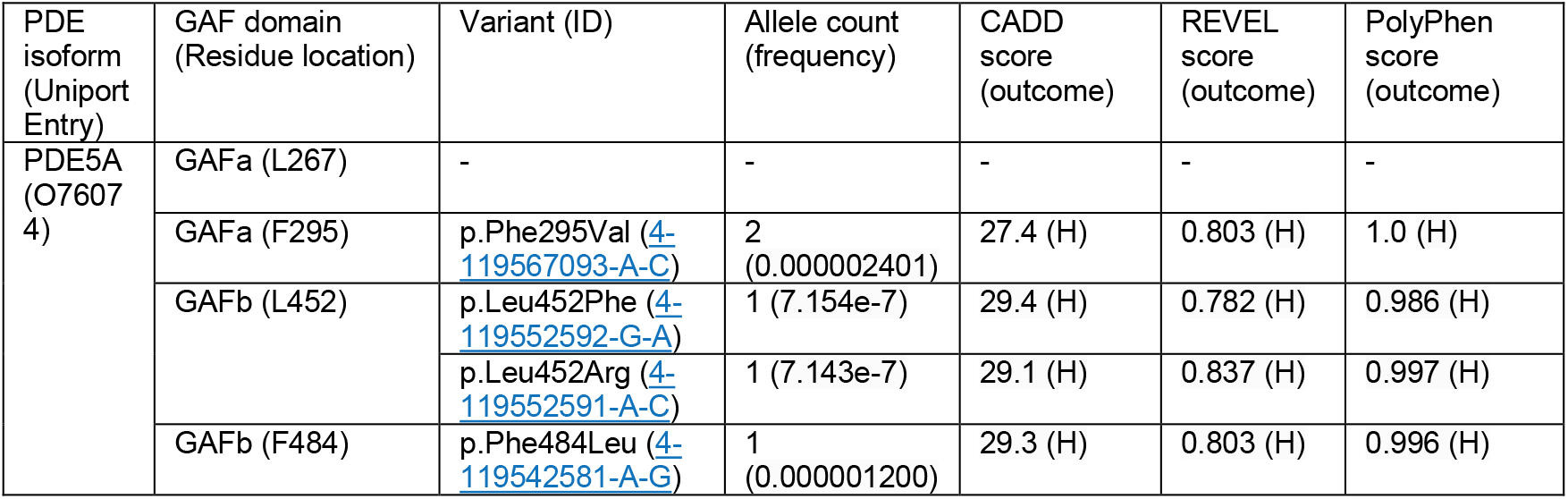
Missense, insertion, and deletion variants of the two coevolving positions in the GAF domain-containing human PDE5 reported in the gnomAD database [75]. CADD [76], REVEL [77], and PolyPhen [78] prediction scores were used to distinguish potentially harmful variants from benign ones. B, potentially benign; H, potentially harmful.

| PDE isoform (Uniport Entry) | GAF domain (Residue location) | Variant (ID) | Allele count (frequency) | CADD score (outcome) | REVEL score (outcome) | PolyPhen score (outcome) |
| --- | --- | --- | --- | --- | --- | --- |
| PDE5A (O76074) | GAFa (L267) | - | - | - | - | - |
|  | GAFa (F295) | p.Phe295Val (4-119567093-A-C) | 2 (0.000002401) | 27.4 (H) | 0.803 (H) | 1.0 (H) |
|  | GAFb (L452) | p.Leu452Phe (4-119552592-G-A) | 1 (7.154e-7) | 29.4 (H) | 0.782 (H) | 0.986 (H) |
|  |  | p.Leu452Arg (4-119552591-A-C) | 1 (7.143e-7) | 29.1 (H) | 0.837 (H) | 0.997 (H) |
|  | GAFb (F484) | p.Phe484Leu (4-119542581-A-G) | 1 (0.000001200) | 29.3 (H) | 0.803 (H) | 0.996 (H) |

**Supporting Table 2:** Predicted effects of the mutations on the stability, flexibility, and functionality of the GAFa domain.

|  | GAFa(L267A) |  | GAFa(F295A) |  |
| --- | --- | --- | --- | --- |
| Tool | Value | Predicted outcome | value | Predicted outcome |
| $\Delta\Delta G$ DynaMut | -3.732 kcal/mol | Destabilizing | -2.172 kcal/mol | Destabilizing |
| $\Delta\Delta G$ mCSM | -3.066 kcal/mol | Destabilizing | -2.867 kcal/mol | Destabilizing |
| $\Delta\Delta G$ SDM | -3.89 kcal/mol | Destabilizing | -3.95 kcal/mol | Destabilizing |
| $\Delta\Delta G$ DUET | -3.716 kcal/mol | Destabilizing | -3.246 kcal/mol | Destabilizing |
| $\Delta\Delta G$ DeepDDG | -3.424 kcal/mol | Destabilizing | -3.504 kcal/mol | Destabilizing |
| $\Delta\Delta G$ iDeepDDG | -3.841 kcal/mol | Destabilizing | -3.452 kcal/mol | Destabilizing |
| $\Delta\Delta G$ Maestro | -2.982 kcal/mol | Destabilizing | -3.183 kcal/mol | Destabilizing |
| $\Delta\Delta G$ PoPMuSiC | -2.28 kcal/mol | Destabilizing | -3.69 kcal/mol | Destabilizing |
| $\Delta\Delta G$ I-Mutant 2.0 | -3.94 kcal/mol | Destabilizing | -3.74 kcal/mol | Destabilizing |
| $\Delta\Delta G$ CUPSAT | -4.85 kcal/mol | Destabilizing | - 3.31 kcal/mol | Destabilizing |
| $\Delta\Delta G$ Mupro | -1.957 kcal/mol | Destabilizing | -1.183 kcal/mol | Destabilizing |
| $\Delta\Delta G$ ENCoM | -0.582 kcal/mol | Destabilizing | -1.321 kcal/mol | Destabilizing |
| $\Delta\Delta S_{vib}$ ENCoM | 0.728 kcal.mol<sup>-1</sup>.K<sup>-1</sup> | Increase flexibility | 1.652 kcal.mol<sup>-1</sup>.K<sup>-1</sup> | Increase flexibility |
| SNAP2 | +59 | Functionally deleterious | +82 | Functionally deleterious |

**Supporting Table 3:** Number of contacts formed by the distant coevolving residue positions in the WT and mutant PDE5 GAFa domain.

| Interaction | WT<br>(L267) | WT<br>(F295) | L267A | F295A |
| --- | --- | --- | --- | --- |
| Polar contacts | 2 | 0 | 2 | 0 |
| Weak polar contacts | 2 | 1 | 3 | 0 |
| Hydrogen bonds | 2 | 0 | 2 | 0 |
| Weak hydrogen bonds | 2 | 1 | 1 | 0 |
| Carbonyl interactions | 1 | 0 | 1 | 0 |
| Hydrophobic contacts | 24 | 25 | 5 | 2 |

### Supporting Figures

**Supporting Figure 1.**
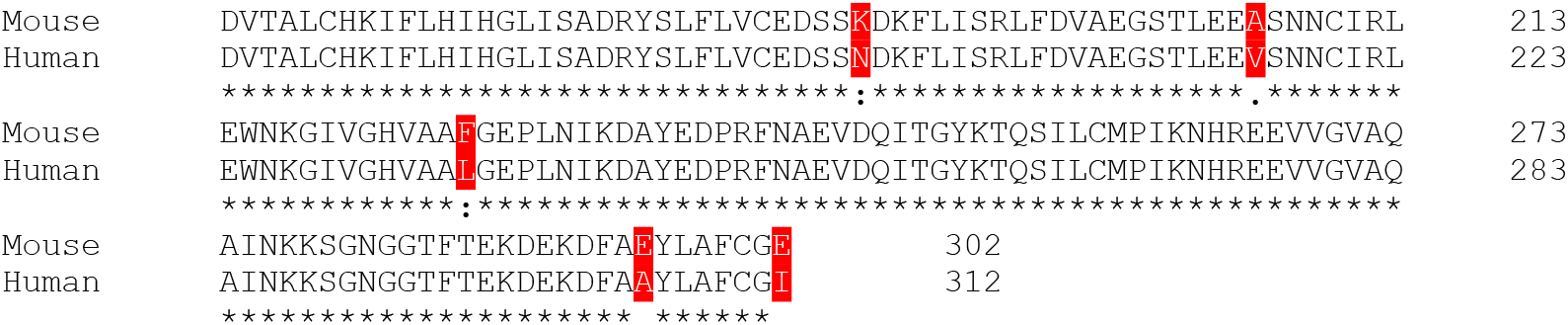
Amino acid sequence alignment of mouse and human PDE5 GAFa domain. Differences in the amino acid sequences are highlighted in red.

**Supporting Figure 2.**
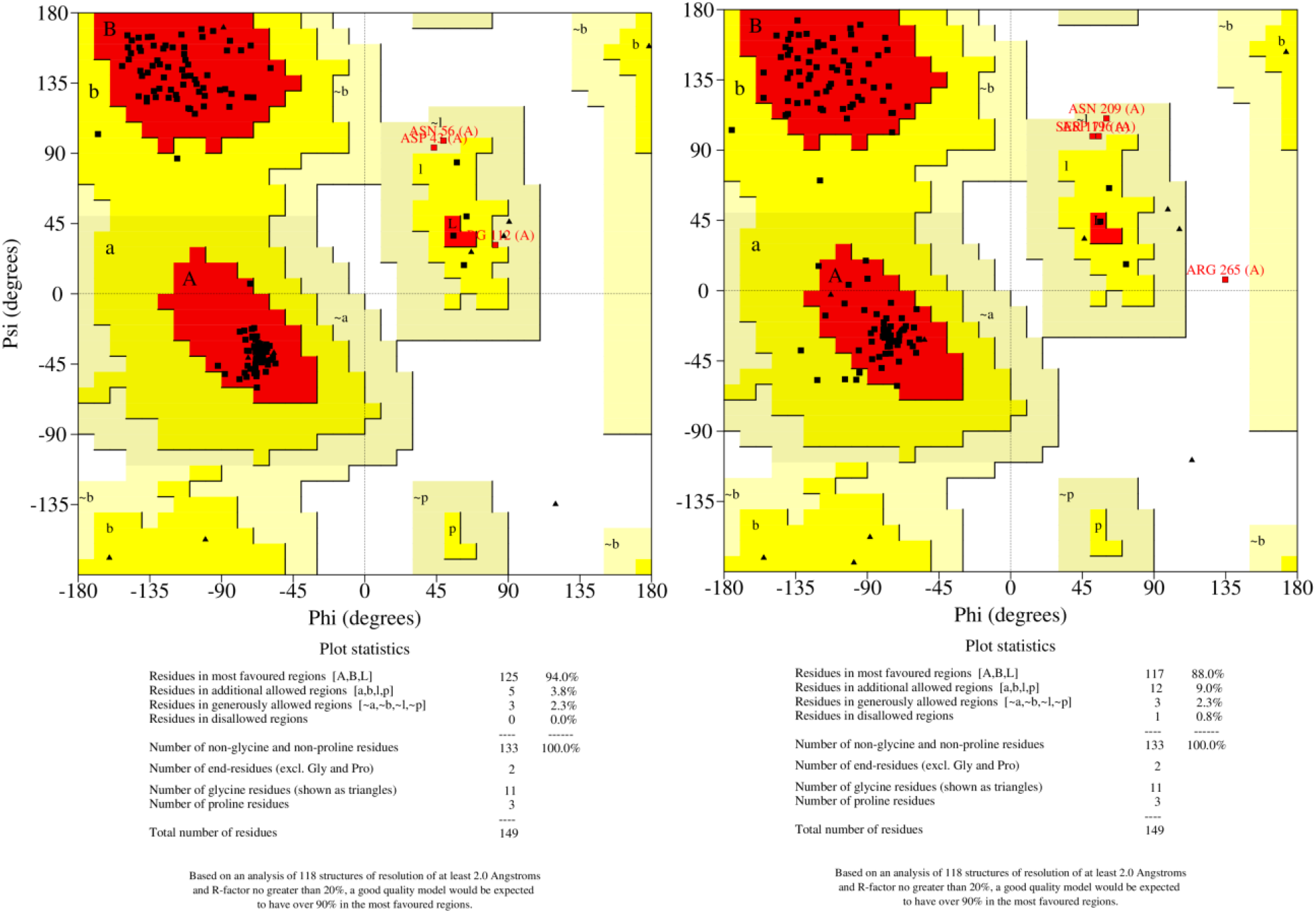
Ramachandran plot of the modeled holo GAFa domain of human PDE5. The structural model of the human GAFa domain of PDE5 (left panel) spanning residues from D164 to I312 was generated from the mouse holo PDE5-GAFa model (PDB: 2K31) (right panel). The core regions (red) represent the most favorable combination of phi-psi values. The percentage of residues in the core is 94%, depicting a better stereochemical quality of the modeled structure compared to the mouse template structure, which has 88% of residues in the core region.

**Supporting Figure 3.**
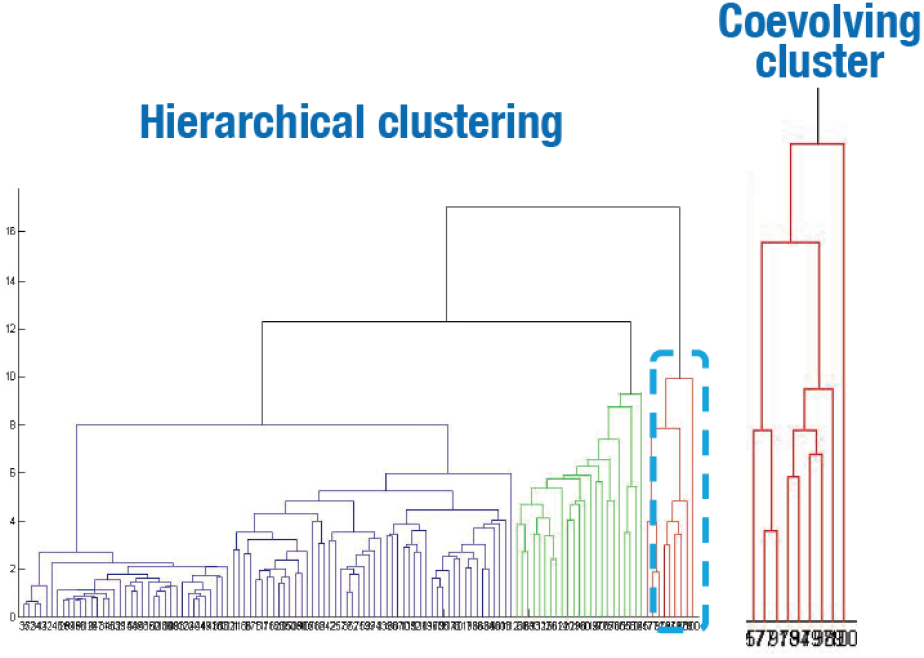
Hierarchical clustering of the coevolving residue positions in the GAF domains based on their pairwise SCA scores. The cluster of residues with the highest SCA scores is shown in red and zoomed in (right panel).

**Supporting Figure 4.**
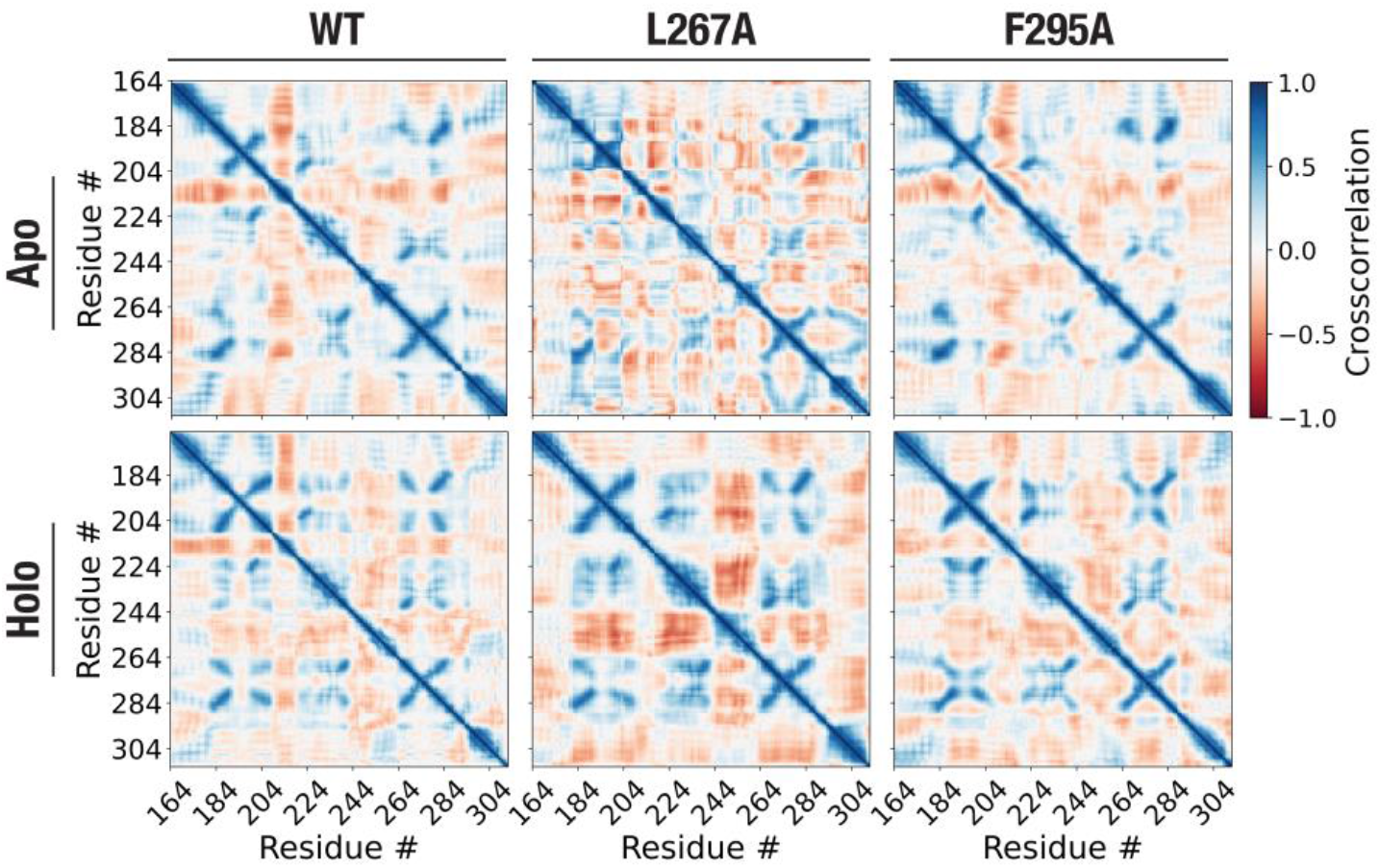
Dynamic cross-correlation (DCC) analysis of apo and holo GAFa domains of human PDE5. Heatmap showing average DCC values of the apo (upper panel) and holo (lower panel) GAFa domains (WT and mutants) obtained from three independent, all-atom, 1000 ns MD simulation runs.

**Supporting Figure 5.**
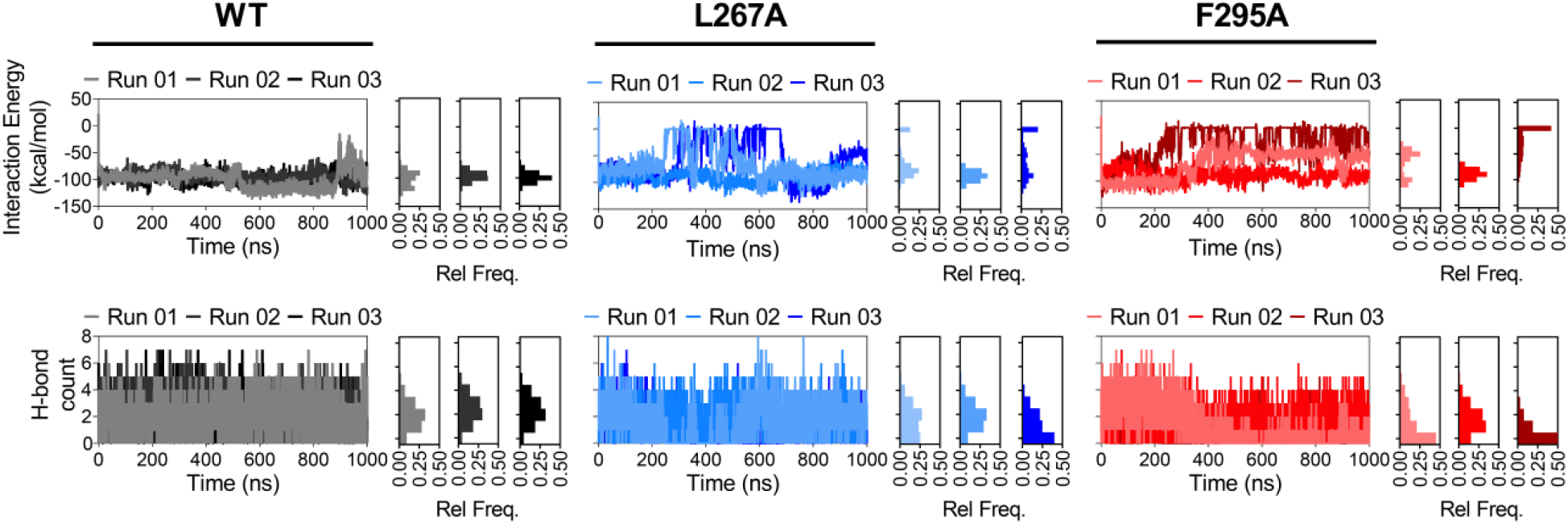
L267A and F295A mutation weakens GAFa domain-cGMP interaction. Graphs showing changes in the interaction energy (vdW and electrostatics; upper panel) and number of H-bonds formed between cGMP and GAFa domain (lower panel) obtained from three independent, all-atom, 1000 ns MD simulation runs. Outsets represent the frequency distribution of interaction energy (upper panel) and H-bonds (bottom panel) obtained from three independent, all-atom, 1000 ns MD simulation runs.

**Supporting Figure 6.**
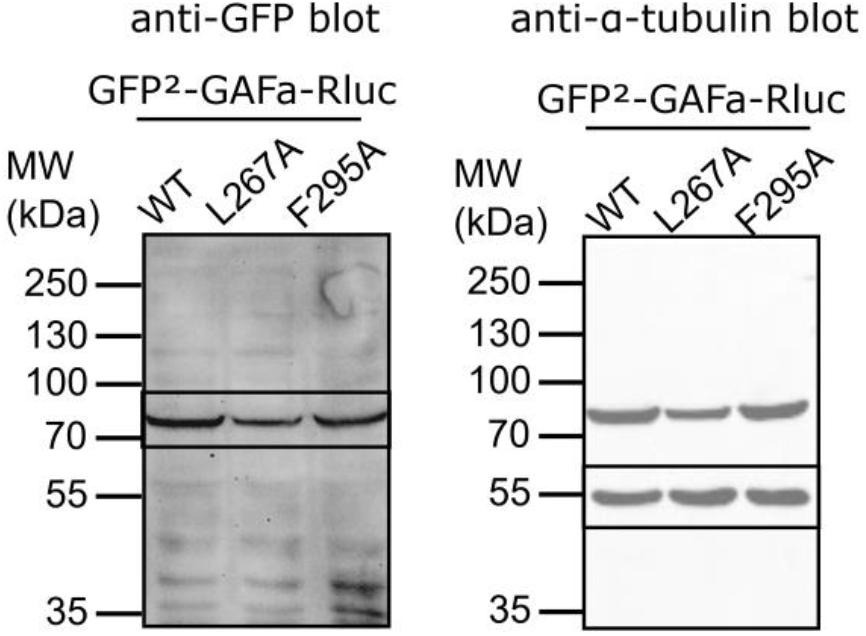
Image of the whole blot used for generating Fig. 4D in the main text. The predicted molecular weight of the GAFa domain biosensor is ∼87 kDa. WT, wild type. Note the boxes in the blot showing the region used to generate the main text figure.

**Supporting Figure 7.**
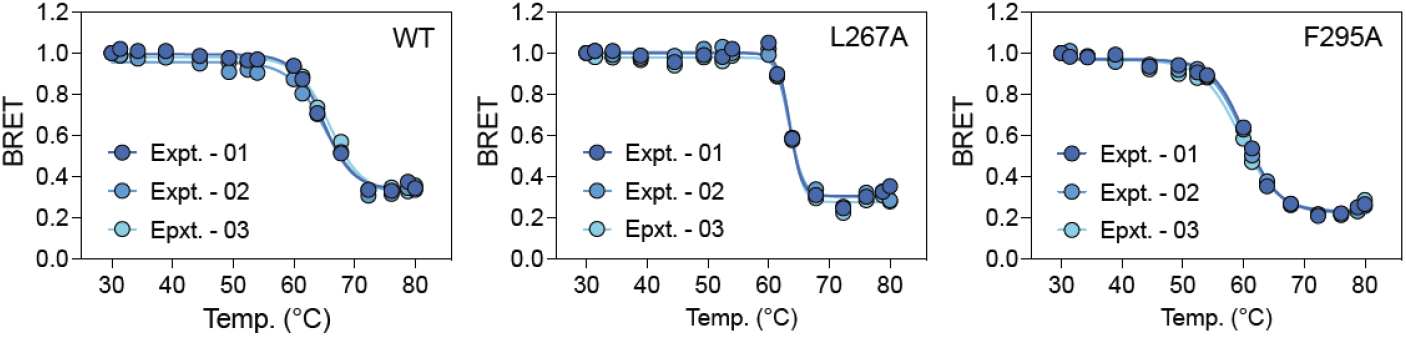
Thermal stability of the GAFa domain. Graph showing the effect of temperature on the stability of the WT and mutant GAFa domains inferred from measuring the BRET ratio between mNG and NLuc in the WT (left) and mutant (middle and right) mNG-GAFa-NLuc biosensors. Data shown are from three independent experiments and fitted to a Boltzmann sigmoidal model to determine melting temperatures (*T*_m_) for each protein.

**Supporting Figure 8.**
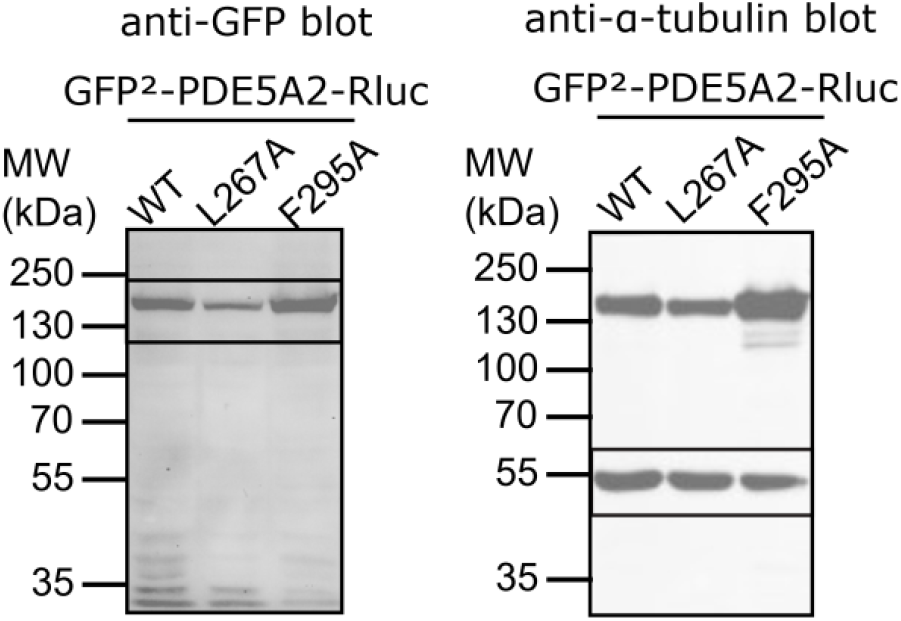
Image of the whole blot used for generating Fig. 5B in the main text. The predicted molecular weight of the full-length PDE5A2 biosensor is ∼162 kDa. WT, wild type. Note the boxes in the blot showing the region used to generate the main text figure.

**Supporting Figure 9.**
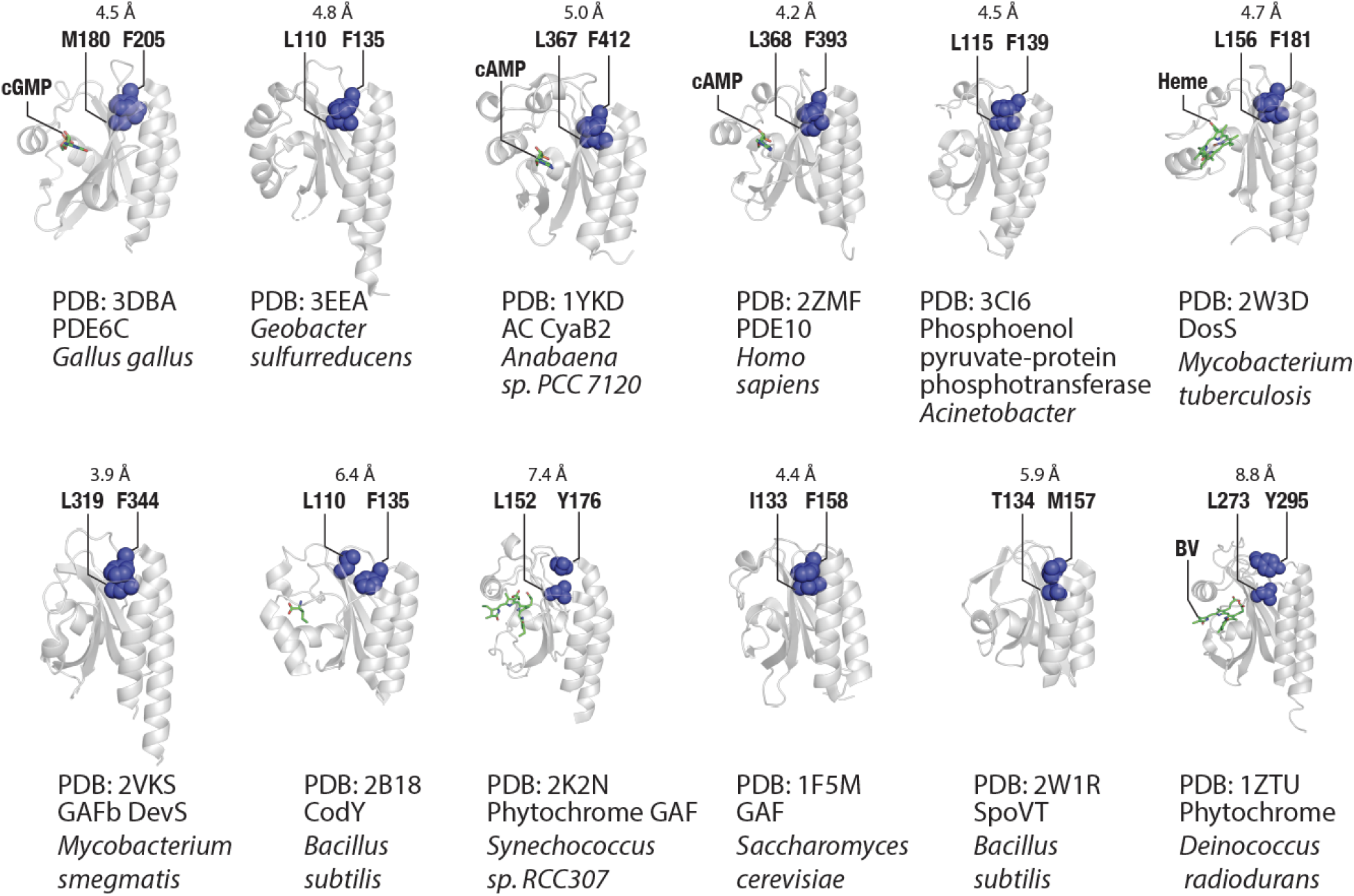
Cartoon representations of GAF domain structures available in the RCSB database including their PDB code, the protein name, and the species name. The ligand, if found in the structure, is represented as sticks, and the distant coevolving residue positions (equivalent to L267 and F295 in the PDE5 GAFa domain) as dark blue spheres. Note the conservation and proximity of the two distant coevolving residue positions in the structures irrespective of the presence of the ligand in the structures.

**Supporting Figure 10.**
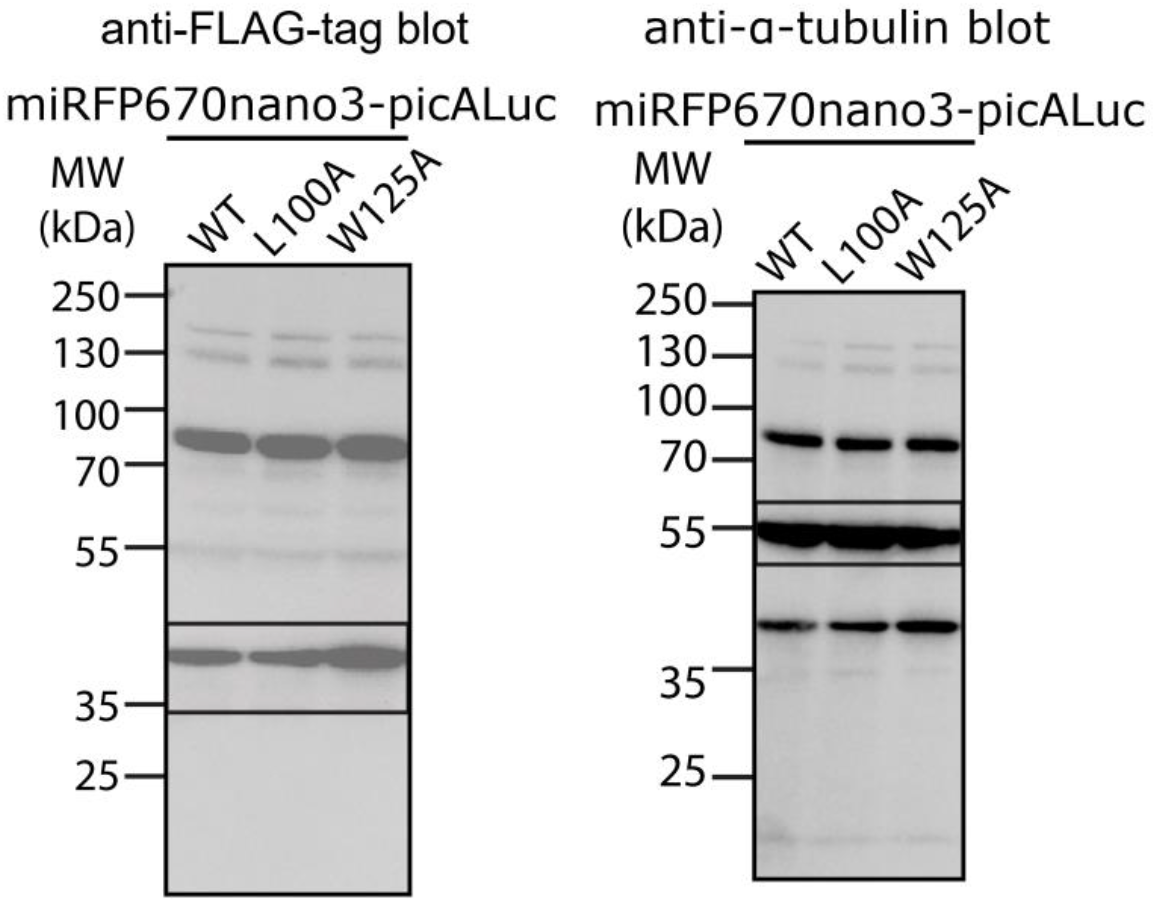
Image of the whole blot used for generating Fig. 5I in the main text. The predicted molecular weight of the miRFP3670nano3 construct is ∼38 kDa. WT, wild type. Note the boxes in the blot showing the region used to generate the main text figure.

**Supporting Figure 11.**
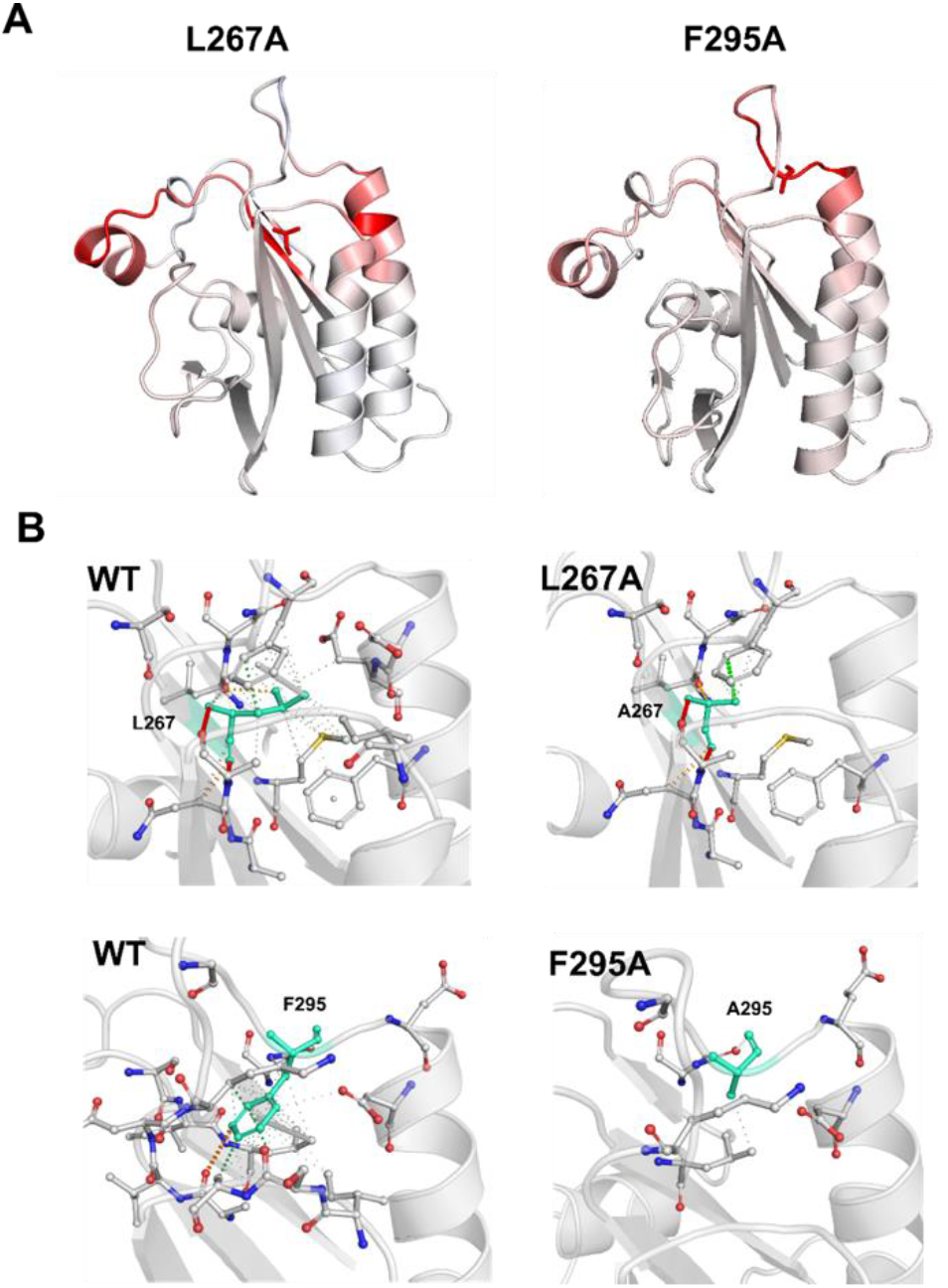
Mutating the coevolving residues disrupts their inter-residue interactions. (A) Change in the vibrational entropy energy (ΔΔSvib) between WT and mutant GAFa domains. Red and blue colors represent gains in flexibility and rigidity, respectively. Mutated residues are shown in red sticks. (B) Inter-residue interactions formed by the mutated positions in the WT (left) and mutant (right) GAFa domains. WT and mutant residues are colored in cyan and represented as sticks alongside the surrounding interacting residues.

### Supporting Movies

**Supporting Movie 1**. Positional mapping of coevolving cluster of residues on the holo GAFa domain of PDE5 (PDB: 2K31).

**Supporting movie 2**. Dynamics of L267 and F295 obtained from the NMR models of PDE5 holo GAFa domain (PDB: 2K31).

**Supporting movie 3**. Comparison of L267 and F295 dynamics in the holo (blue spheres) and apo (purple spheres) GAFa domain of PDE5.

**Supporting movie 4**. MD simulation - Apo WT GAFa domain Run 01

**Supporting movie 5**. MD simulation – Apo WT GAFa domain Run 02

**Supporting movie 6**. MD simulation – Apo WT GAFa domain Run 03

**Supporting movie 7**. MD simulation – Apo L267A GAFa domain Run 01

**Supporting movie 8**. MD simulation – Apo L267A GAFa domain Run 02

**Supporting movie 9**. MD simulation – Apo L267A GAFa domain Run 03

**Supporting movie 10**. MD simulation – Apo F295A GAFa domain Run 01

**Supporting movie 11**. MD simulation – Apo F295A GAFa domain Run 02

**Supporting movie 12**. MD simulation – Apo F295A GAFa domain Run 03

**Supporting movie 13**. MD simulation - Holo WT GAFa domain Run 01

**Supporting movie 14**. MD simulation – Holo WT GAFa domain Run 02

**Supporting movie 15**. MD simulation – Holo WT GAFa domain Run 03

**Supporting movie 16**. MD simulation – Holo L267A GAFa domain Run 01

**Supporting movie 17**. MD simulation – Holo L267A GAFa domain Run 02

**Supporting movie 18**. MD simulation – Holo L267A GAFa domain Run 03

**Supporting movie 19**. MD simulation – Holo F295A GAFa domain Run 01

**Supporting movie 20**. MD simulation – Holo F295A GAFa domain Run 02

**Supporting movie 21**. MD simulation – Holo F295A GAFa domain Run 03

